# Oligomeric HIV-1 Integrase Structures Reveal Functional Plasticity for Intasome Assembly and RNA Binding

**DOI:** 10.1101/2024.01.26.577436

**Authors:** Tao Jing, Zelin Shan, Tung Dinh, Avik Biswas, Sooin Jang, Juliet Greenwood, Min Li, Zeyuan Zhang, Gennavieve Gray, Hye Jeong Shin, Bo Zhou, Dario Passos, Timothy S. Strutzenberg, Sriram Aiyer, Leonardo Andrade, Yuxuan Zhang, Zhen Li, Robert Craigie, Alan N. Engelman, Mamuka Kvaratskhelia, Dmitry Lyumkis

**Author notes:** Correspondence: Dmitry Lyumkis. These authors contributed equally to this work (Tao Jing, Zelin Shan).

## Abstract

Integrase (IN) performs dual essential roles during HIV-1 replication. During ingress, IN functions within an oligomeric “intasome” assembly to catalyze viral DNA integration into host chromatin. During late stages of infection, tetrameric IN binds viral RNA and orchestrates the condensation of ribonucleoprotein complexes into the capsid core. The molecular architectures of HIV-1 IN assemblies that mediate these distinct events remain unknown. Furthermore, the tetramer is an important antiviral target for allosteric IN inhibitors. Here, we determined cryo-EM structures of wildtype HIV-1 IN tetramers and intasome hexadecamers. Our structures unveil a remarkable plasticity that leverages IN C-terminal domains and abutting linkers to assemble functionally distinct oligomeric forms. Alteration of a newly recognized conserved interface revealed that both IN functions track with tetramerization *in vitro* and during HIV-1 infection. Collectively, our findings reveal how IN plasticity orchestrates its diverse molecular functions, suggest a working model for IN-viral RNA binding, and provide atomic blueprints for allosteric IN inhibitor development.

## Introduction

Human immunodeficiency virus 1 (HIV-1) currently infects ∼40 million people worldwide, and the number of infected individuals continues to rise. Left alone, HIV-1 will eventually lead to acquired immunodeficiency syndrome through a series of progressive events that weaken the immune system and make individuals susceptible to opportunistic infections. Although there are now numerous therapies for people living with HIV that block different stages of the viral replication cycle, HIV-1 infection afflicts individuals for life, and there remains no cure^1^.

After cell entry, HIV-1 RNA is reverse transcribed into double-stranded linear DNA containing a copy of the viral long terminal repeat (LTR) at each end. A hallmark of HIV-1 infection is the covalent insertion of this viral DNA (vDNA) into host chromatin, the terminal step in viral ingress that is necessary to establish a persistent infection^2,3^. This key step, termed integration, distinguishes HIV-1 and related retroviruses from other animal viruses and poses a central challenge to therapeutic intervention. Concerted integration of both LTR ends catalyzed by IN^4^ requires the formation of a nucleoprotein complex containing multimers of IN bound to the vDNA ends, termed “intasomes”^5^. Intasomes can be assembled *in vitro* from purified IN and vDNA oligonucleotides and are catalytically competent^6,7^. Structural biology efforts elucidated how a conserved HIV-1 intasome core consisting of two IN dimers scaffolds and intertwines the vDNA ends^8^. However, that same study revealed a putative larger intasome architecture, consisting of up to sixteen protomers of IN, which could not be fully resolved or modeled into the experimental density^8^. Structures of related lentiviral intasomes from Maedi Visna virus (MVV)^9,10^, which are more stable and yield homogeneous assemblies, have suggested that lentiviral intasomes might be generally hexadecameric, i.e. arise from sixteen copies of IN^11^. However, all HIV-1 intasome structures resolved to date^8,12–16^ lack key IN flanking subunits that assemble on the intasome periphery, and atomic models constitute only ∼40% of the total protein mass. These limitations can be attributed to the dynamic nature of HIV-1 IN^17,18^ (also observed with simian immunodeficiency virus IN^19^), characteristics that confer significant challenges to structural biology attempts. Thus, whether the hexadecameric intasome architecture is conserved in HIV-1 remains unclear.

Viral genetics initially revealed pleiotropic outcomes of HIV-1 IN mutagenesis. Some substitutions, notably those that affect active site residues (i.e., Asp64, Asp116, and Glu152), could specifically block integration in infected cells, consistent with selective abrogation of intasome function^20–23^. These were subsequently termed class I IN mutations^24^. In contrast, the majority of IN substitutions impair additional processes, including reverse transcription as well as particle assembly and release from transfected cells (reviewed in reference^24^). These were termed class II IN mutations^24^. The presence of multiple classes of mutations affecting distinct stages of viral replication highlighted the complexity of IN and suggested multifaceted functionalities during HIV-1 infection^25^.

Recent work has shown that the compendium of class II substitutions can be explained by a second essential function of IN that exerts itself during the late stage of viral replication. HIV- 1 IN binds directly to viral RNA (vRNA) in virions, and impairing the IN-vRNA interaction yields eccentric, non-infectious virions characterized by mis-localization of ribonucleoprotein complexes (RNPs) outside of the protective capsid core^26^. Multiple lines of evidence emerged indicating the essential role of IN-vRNA interactions for virion maturation: i) allosteric IN inhibitors (ALLINIs) impaired the binding of wildtype (WT) IN, but not the drug-escape IN A128T mutant, to vRNA in HIV-1 virions^26^; ii) class II R269A/K273A substitutions compromised IN binding to vRNA in virions without significantly affecting other known IN functions and resulted in non-infectious particles with eccentrically mislocalized RNPs^26^; and iii) viral evolution studies have revealed the emergence of compensatory mutations within the IN C-terminal domain (CTD) that specifically restored IN interactions with vRNA^27^. Biochemical assays suggested that IN tetramers, but not dimers or monomers, effectively bind cognate vRNA segments, as amino acid substitutions that impaired IN tetramerization also inhibited IN-vRNA binding *in vitro* and in virions^28^. Collectively, these results proposed a novel biological role for IN-vRNA binding during virion morphogenesis^26,28^. However, the structural organization of the IN tetramer, which mediates vRNA binding, remains unknown.

Furthermore, IN tetramers are the antiviral targets for investigational ALLINIs^29^, which are also known by alternative names such as LEDGF-IN site inhibitors (LEDGINs)^30^, IN-LEDGF allosteric inhibitors (INLAIs)^31,32^, multimerization IN inhibitors (MINIs)^33^, or noncatalytic IN inhibitors (NCINIs)^34,35^. Of note, the ALLINI pirmitegravir is currently in Phase 2 clinical trial (Identifier: NCT05869643). ALLINIs potently impair proper virion maturation by inducing aberrant or hyper multimerization of IN, which inhibits vRNA binding^26,29,36,37^. Consequently, the virions produced in the presence of ALLINIs have RNPs mis-localized outside of the mature capsid and are non-infectious. Previous structural studies have elucidated the mode of action of ALLINIs in the context of full-length dimeric IN mutant Y15A/F185H, which is substantially resistant to ALLINIs^38^, or artificial constructs that accentuate intermolecular IN-IN interactions to template aberrant IN hyper multimerization^39^. The authentic antiviral target of these inhibitors, however, is the full-length WT IN tetramer^29^. Therefore, elucidating the structure of an IN tetramer is of significant pharmacological importance.

HIV-1 IN consists of three functional domains including the N-terminal domain (NTD), catalytic core domain (CCD), and CTD^40^. The NTD harbors a zinc-binding motif that stabilizes its fold and helps coordinate the assembly of the multimeric intasome complex^41^. The CCD harbors the conserved DDE motif containing the Asp64-Asp116-Glu152 catalytic triad^42^ and is involved in the cleavage and joining of vDNA strands for integration. The CTD plays an essential role in DNA binding, IN multimerization, and the proper positioning of IN on the vDNA ends to promote the assembly of the higher-order intasome^6,8,43^. Notably, electropositive residues within the CTD primarily mediate IN-vRNA binding^26,28^. All three domains are required for IN to properly execute its two distinct functions. However, despite nearly three decades of structural biology work that contributed to our growing understanding of IN function (reviewed in ^2^), structures of architecturally complete HIV-1 IN assemblies – whereby each of the three domains constituting all protomers within the complex can be defined – have remained critically absent.

Here, we used cryo-EM to determine structures of two oligomeric assemblies formed by WT full-length HIV-1 IN. First, using the scaffolding domain from an IN host factor to stabilize and solubilize tetrameric IN, we present the structure of the IN tetramer, which is characterized by interlocked placement of all four CTDs that maintain the architectural integrity of the tetrameric assembly. Next, using a biochemical strategy to isolate stabilized cross-linked intasomes and focused classification approaches during image analysis of cryo-EM data, we deciphered the architectural assembly of the hexadecameric WT HIV-1 intasome, which revealed how coordinated structural rearrangements that arise within the IN tetramer facilitate assembly into the complete intasome. Inspection of the contacts mediated by the NTD-CTD interactions revealed two conserved residues, E35 and K240, which form a salt bridge that contributes to both the structural integrity of an IN tetramer and the interactions observed within the hexadecameric intasome. Our reverse-charge guided mutagenesis studies targeting these interactions have revealed that both catalytic and non-catalytic functions of IN *in vitro* and in infected cells closely correlated with the ability of the mutant proteins to form tetramers. Using molecular modeling, supported by IN mutagenesis, multimerization analyses, and RNA bridging assays, we suggest how vRNA may bind the IN tetramer. These data highlight how the plastic molecular architecture of HIV-1 IN can mediate its diverse functions and provide a molecular blueprint for the design of novel inhibitors that can block IN function at distinct stages of the viral replication cycle.

## Results

### Interlocked CTDs maintain the architectural integrity of the IN tetramer

Prior studies suggested that the formation of a core IN tetramer is relevant for the proper execution of two distinct functions mediated by IN, integration^9^ and RNA binding^26–28^. We set out to determine the structure of the WT IN tetramer, without bound nucleic acids. Previous work showed that the IN-binding domain (IBD) of the IN host factor lens epithelium-derived growth factor (LEDGF)/transcriptional co-activator p75 can solubilize and stabilize tetrameric IN^17,18,44,45^. We thus included the IBD in our bacterial co-expression system, which was expected to tightly bind and stabilize a functional oligomeric form of IN to facilitate its structure determination. Size exclusion chromatography (SEC) experiments revealed that in the absence of the IBD, IN predominantly formed dimers and tetramers in the presence and absence of CHAPS detergent, respectively. As anticipated^17,18^, IBD co-expression predominantly yielded IN tetramers (**Supplementary Fig. 1**). The IN-IBD complex was purified using nickel affinity chromatography followed by SEC, yielding a peak corresponding to the appropriate molecular weight of the tetramer and containing both IN and IBD polypeptides (**Supplementary Fig. 2**). We advanced this complex for structural biology studies. The IN-IBD particles exhibited severe preferential orientation on cryo-EM grids. To address this limitation, we tilted the stage during data collection^46–48^. We collected 2,649 movies of the IN-IBD complex and incorporated iterative 2D and 3D classification in cryoSPARC^49, 50^producing a final map resolved to ∼4.1 Å and an atomic model derived from this map (**Supplementary Fig. 3**-5 and **Supplementary Table 1**). We also collected a large cryo-EM dataset of apo IN in the absence of bound IBD, and reconstructed a map to ∼7 Å resolution (**Supplementary Fig. 6**-7). The resulting structure revealed the same general IN tetramer architecture as the IBD-engaged complex, confirming that the presence of IBD does not grossly alter the structural organization of tetrameric IN (**Supplementary Fig. 8**).

The IN-IBD structure is defined by two-fold rotational symmetry and contains four IN protomers, with one IBD monomer bound to each of the four corners of the tetrameric assembly (**Figure 1a-b**). All three of the functionally relevant IN domains, the NTD, CCD, and CTD, are fully resolved within the density. The NTDs and CCDs are connected by ∼15-residue flexible linkers^25^.

**Figure 1.**
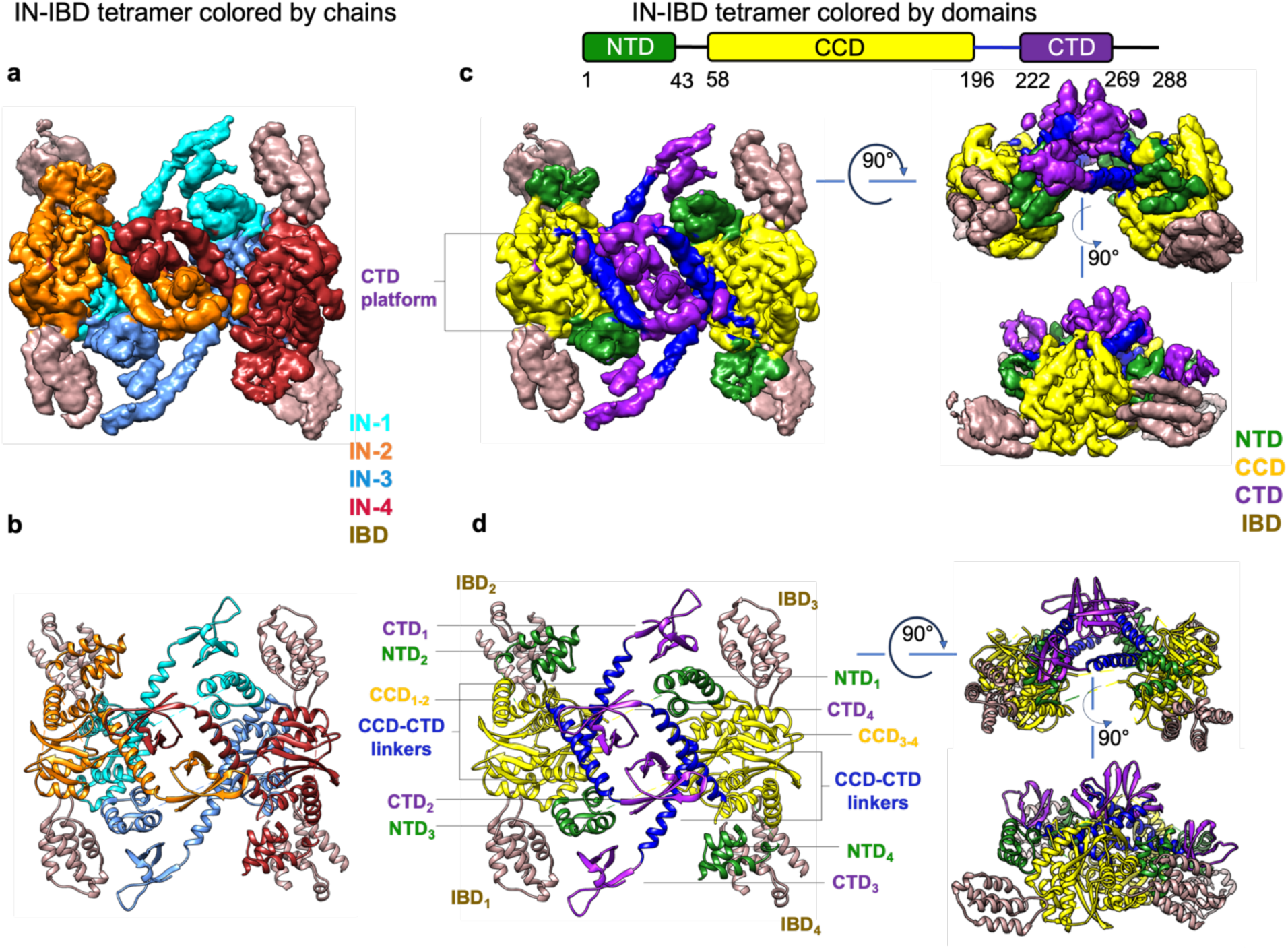
Cryo-EM structure of the tetrameric HIV-1 IN-IBD complex. (**a**) Cryo-EM reconstruction. Colors refer to individual IN protomers (orange, cyan, light blue, red) and IBD (brown). (**b**) Atomic model colored by IN protomers as in **a**. (**c**) Segmented cryo-EM density colored by IN domains (NTD, green; CCD, yellow; CCD-CTD linker, blue; CTD, purple) and IBD (brown). (**d**) Atomic model colored by IN domains as in **c**. Subscripts denote the four different IN and IBD chains.

Accordingly, these linkers were not modeled, although density at low threshold was apparent, which informed their connectivity to the respective CCDs. In contrast, all four linkers connecting the CCDs to the CTDs are alpha-helical and fully defined within the cryo-EM density. As expected, the conserved CCD-CCD interface^51^ scaffolds each of the two IN dimers^6,8^. The tetrameric organization is held together by the organization of the CTDs. The two inner CTDs extend and interlock with one another through the CCD-CTD linkers, whereas the two outer CTDs sandwich the core assembly. The role of the CTD platform in maintaining the integrity of the tetramer is evident when the structures are colored by individual IN domain and viewed by 90° orientations (**Figure 1c-d**). Displayed as such, it becomes apparent that all four CTDs are arranged at the “top” of the assembly, whereas the CCD:CCD dimers reside underneath. All four NTDs are bound to the CCD:CCD dimer interface, with NTD_1_/NTD_3_ emanating from opposing dimers to help maintain the tetrameric configuration. This arrangement was previously observed for two-domain assemblies lacking the CTD^18,52^. The primary binding mode for each IBD resembles prior crystal structures with isolated CCD and NTD-CCD constructs (**Supplementary Fig. 9**a)^53,54^. The binding mode is asymmetric, with density for IBD_1_/IBD_3_ being more pronounced than the density for IBD_2_/IBD_4_ (**Supplementary Fig. 9**b). We attribute this asymmetry, at least in part, to the fact that IBD_1,3_ interface with both NTD_3,1_ and with CTD_3,1_ (weakly), whereas IBD_2,4_ engage only NTD_2,4_. The weak IBD_1,3_–CTD_3,1_ interactions may explain why the IBD helps stabilize and solubilize the tetrameric form of IN^17,18^.

Collectively, our IN-IBD structure reveals how the CTDs bridge two independent IN dimers through alpha-helical linkers. The presence of numerous interfaces explains why all three domains are required for functional IN oligomerization.

### Conformational plasticity within inter-domain linkers leads to context-dependent interfaces made by the CTDs

The CTDs exhibit a highly flexible positional configuration in relation to the dimeric CCD scaffold. Superposition of each IN dimer within the tetrameric IN-IBD assembly to previously determined crystal structures of two-domain HIV-1 IN CCD-CTD constructs (PDBs 1EX4^55^ and 5HOT^38^) or from lentiviruses MVV (PDB 5T3A^10^) or SIV (PDB 1C6V^56^) revealed rearrangements in both position and configuration of the CTDs, mediated by the helical linkers (**Figure 2a**). The most dramatic demonstration of this structural plasticity is observed by comparing the configurations of the CTDs from the HIV-1 IN tetramer with six independent modeled IN protomers from the MVV intasome, which are structurally homologous (PDB 7U32^10^; two of these eight protomers did not contain atomic models of the CTD) (**Figure 2b**). These comparisons imply that the conformational plasticity of the CCD-CTD linker plays a key role in choreographing diverse assemblies, including IN oligomers and the intasome.

**Figure 2.**
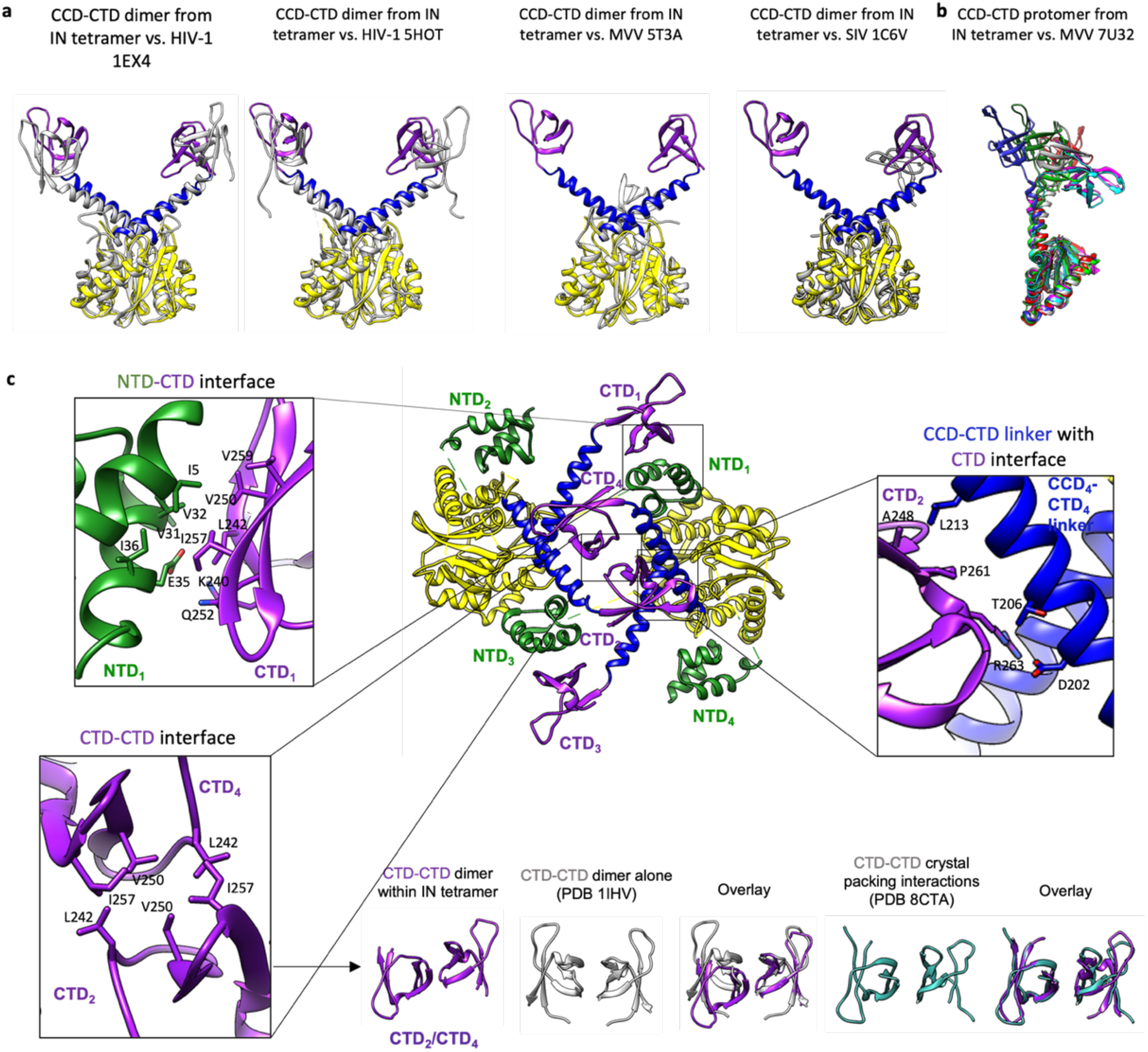
The structure of full-length IN reveals novel conformational rearrangements and inter- domain interactions. (**a**) Comparison of the CCD-CTD dimer from the IN tetramer to previously determined two-domain IN structures from HIV-1 or other lentiviruses. All prior structures are colored in gray and correspond to HIV-1 (1EX4^55^ and 5HOT^38^), MVV (5T3A)^10^, or SIV (1C6V^56^). (**b**) Comparison of a single CCD-CTD protomer from the IN tetramer to individual fully modeled IN protomers from the MVV intasome^10^. The conformation of the CCD-CTD protomer from HIV-1 is distinct from all modeled MVV conformations, implying a large degree of structural plasticity. A single CCD-CTD protomer from the IN tetramer is colored in gray. (**c**) The structure of the tetramer is displayed in the center, with key interaction interfaces shown within inset panels. The interaction mediated by the central CTD:CTD dimer (bottom left panel) resides in an anti-parallel configuration, which is distinct from the parallel configuration previously encountered in CCD-CTD oligomeric assemblies with bound ALLINIs^38^ or within dimeric CTD:CTD structures in solution^57^ (shown in gray, PDB: 1IHV), but resembles the crystal packing interactions observed in a recent ALLINI-bound structure (PDB: 8CTA)^58^ shown in light sea green. The CTD also mediates other diverse interactions, including with the NTD (upper left) and with the CCD-CTD linker (upper right) within the IN tetramer.

As a consequence of the structural plasticity, the CTDs can engage in diverse interactions (**Figure 2c**). There are two prominent interactions between the CTDs and NTDs on the peripheral edges of the tetrameric complex, which are likely relevant for securing the outer CTDs (**Figure 2c**, top left). This interface is also conserved in the HIV-1 intasome, although it was not previously appreciated (**Supplementary Fig. 10**). The IN tetramer is also stabilized by interactions between the CTD of one protomer and the CCD-CTD linker region of another protomer (**Figure 2c**, top right). Lastly, the CTDs engage one another at the twofold axis through hydrophobic contacts (**Figure 2c**, bottom left). This engagement is anti-parallel, which differs from the original dimeric CTD solution NMR structure (PDB 1IHV)^57^, but resembles recently described crystallographic packing interactions within ALLINI-IN assemblies (PDB 8CTA)^58^ (**Figure 2c**, bottom). Notably, the fact that the anti-parallel CTD:CTD interface is observed via cryo-EM within the IN tetramer indicates that this interaction mode is likely biologically relevant, which could not be concluded from packing assemblies observed crystallographically^58^. Such diverse contacts contributed by CTDs help stabilize and maintain the tetrameric conformation and add to our understanding of the promiscuity of the CTD, which additionally engages with other protein domains and/or nucleic acids in the intasome^8^.

In contrast to the promiscuous CTD interactome, the positions of the NTDs are more conserved. The NTDs are typically bound to the two-fold interface of each CCD dimer, a conserved location among IN structures, including intasome assemblies (**Supplementary Fig. 11**). The NTDs can rearrange between protomers. For example, in the tetramer structure, NTD_1_/NTD_3_ extend out across opposing dimers; also, the NTDs rearrange upon intasome assembly, which is required for intertwining the vDNA. In all cases, however, the resulting CCD:NTD interfaces are maintained^6–8,10,59–62^. The CCDs are universally conserved and engage through the canonical dimerization interface^51^.

These data highlight the propensity of the CTD to form distinct context-dependent interactions, which explains why this domain is key to higher-order IN assembly and mediates its functional plasticity.

### The IN tetramer forms the core organizational unit of the hexadecameric HIV-1 intasome

Prior lentiviral intasome structures have revealed a diversity of oligomeric IN species ranging from tetramer to hexadecamer^8–10,12,13,15,19^, and recent results suggested that functional IN tetramerization is essential for lentiviral intasome assembly^9^. However, owing to the propensity of HIV-1 IN to aggregate *in vitro*, and to challenges with biochemical intasome assembly, structures of HIV-1 intasomes containing fully resolved flanking subunits have remained elusive. This limitation has precluded an understanding of how HIV-1 IN assembles into the complete intasome.

To address this limitation, we set out to determine the complete structure of the WT HIV- 1 intasome and resolve the missing flanking regions. Intasomes were assembled using a similar strategy to those previously described^8,12,13^, but we additionally used the drug Dolutegravir to prevent spurious strand transfer^63^ and crosslinked the sample to stabilize discrete multimeric forms and isolate assemblies only within the expected ∼600 kDa range (**Supplementary Fig. 12**). The eluted mixture was vitrified on grids and subjected to cryo-EM analysis. We collected a dataset consisting of 775 movies and used standard protocols for image refinement and classification, yielding a map resolved to 3.5 Å within the intasome core. However, despite the crosslinking and extensive biochemical purification, density for the flanking regions remained unresolved (**Supplementary Fig. 13**). We thus sought to recover the densities computationally. We subjected the dataset to multiple rounds of global classification, followed by density subtraction and focused classification (**Supplementary Fig. 14**). Using an iterative approach, we could resolve each half of the intasome separately and use rigid body docking to fit all constituent domains. Importantly, independent analysis of both halves of the intasome yielded nearly identical domain docking, indicating, as expected, that the architectural integrity and mode of IN oligomerization is conserved across each half of the two-fold symmetric complex. We could then derive a composite map of the hexadecameric HIV-1 intasome, assembled from two independently refined halves (**Supplementary Fig. 15**). Using rigid-body docking, we assembled a pseudo-atomic model from the composite map (**Supplementary Fig. 16**).

Four IN tetramers comprise the hexadecameric HIV-1 intasome (**Figure 3a**). The two-fold symmetry of the intasome makes tetramers I/II and III/IV identical. Each of the two unique tetramers have two CCD:CCD dimers that are juxtaposed about a central set of interconnected CTDs, with the CTD-CCD linkers interlocking the arrangement into an architecture that is similar to that observed for the tetrameric IN-IBD structure. Notably, the conformational flexibility of CTD-CCD linkers facilitates the incorporation of each tetramer into the intasome hexadecamer. The major structural rearrangements involve the “compression” of each tetramer about the interlocked CTDs, bringing the two dimers into closer proximity when compared to the apo tetramer (**Figure 3b-c**). Accompanying changes include small repositioning of the CTDs residing on the inner CCD protomers, which in turn interface with other regions of the intasome, as well as a well-known and conserved rearrangement of the NTDs around the vDNAs^2,3^ (depicted by arrows). As expected, the HIV-1 intasome has an overall similar architectural arrangement and protomer organization as the MVV intasome, albeit with weaker interactions maintaining tetramer-tetramer contacts through bridging CTDs (**Supplementary Fig. 17**). Guided by prominent experimental densities for alpha helical linker assignments, we could also fully assign individual domains for each of the sixteen constituent protomers, thus establishing the complete hexadecameric HIV-1 intasome architecture. Collectively, these results highlight complex structural rearrangements, spanning individual IN domains and quaternary assemblies, that must arise to ensure proper intasome formation.

**Figure 3.**
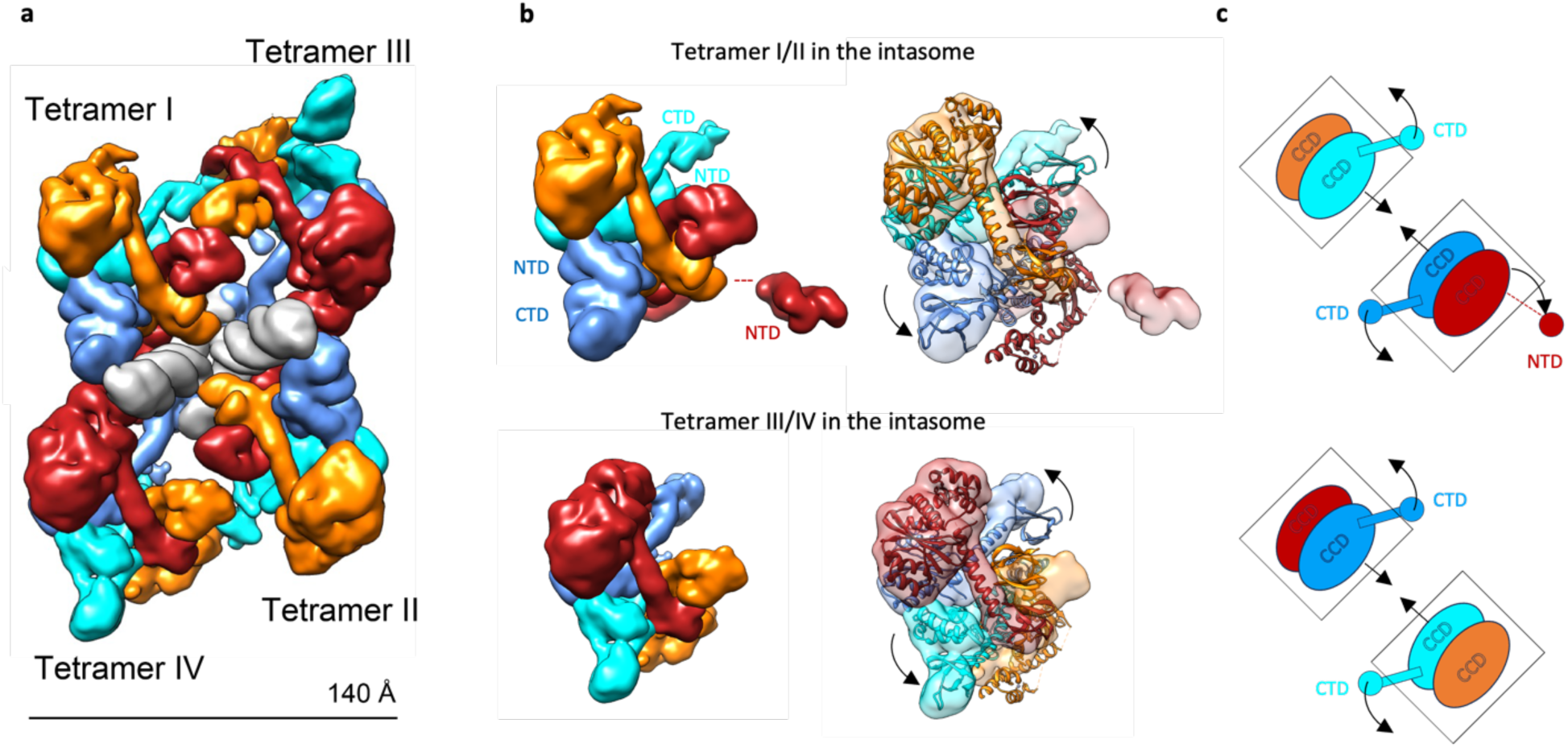
The IN tetramer undergoes compression and domain rearrangements to assemble the hexadecameric HIV-1 intasome. (**a**) Composite map of the complete HIV-1 intasome with four IN tetramers assembled with two vDNA ends. Each of four IN protomers are colored in red, orange, blue, or cyan; the vDNA is in gray. (**b**) Two distinct IN tetramers from the hexadecameric HIV-1 intasome are displayed as isosurfaces (left) and with the apo IN tetramer docked into the density (right). The two IN dimers constituting the apo tetramer must move closer together, or compress, to fit into each of the tetrameric building blocks within the HIV-1 intasome. This is apparent from the bottom dimer, which resides outside of the EM density, whereas the top dimer nearly perfectly superimposes. The curved arrows indicate rearrangements of NTDs and CTDs. (**c**) Schematic showing all major structural changes of the HIV-1 IN tetramer to accommodate intasome assembly. Boxes around the CCDs highlight the compression of individual IN dimers, whereas the curved arrows indicate rearrangements of NTDs and CTDs.

### Functional implications of the tetrameric assembly in vitro and in infected cells

IN tetramerization is thought to play an important role both in the early and the late stages of HIV-1 replication^9,18,26,28,29^. To critically test this model, we searched for likely stabilizing interactions within the apo IN tetramer and identified a salt bridge between E35 of the NTD and K240 of the CTD on the peripheral edges of the complex (**Figure 2c** and **4a**). This interaction is conserved in the HIV-1 intasome (**Supplementary Fig. 10**), although its functional relevance was previously untested. We posited that the interaction between E35 and K240 may in fact be relevant to the biochemical and functional properties of IN, and we sought to interrogate the relevance of this interaction.

**Figure 4.**
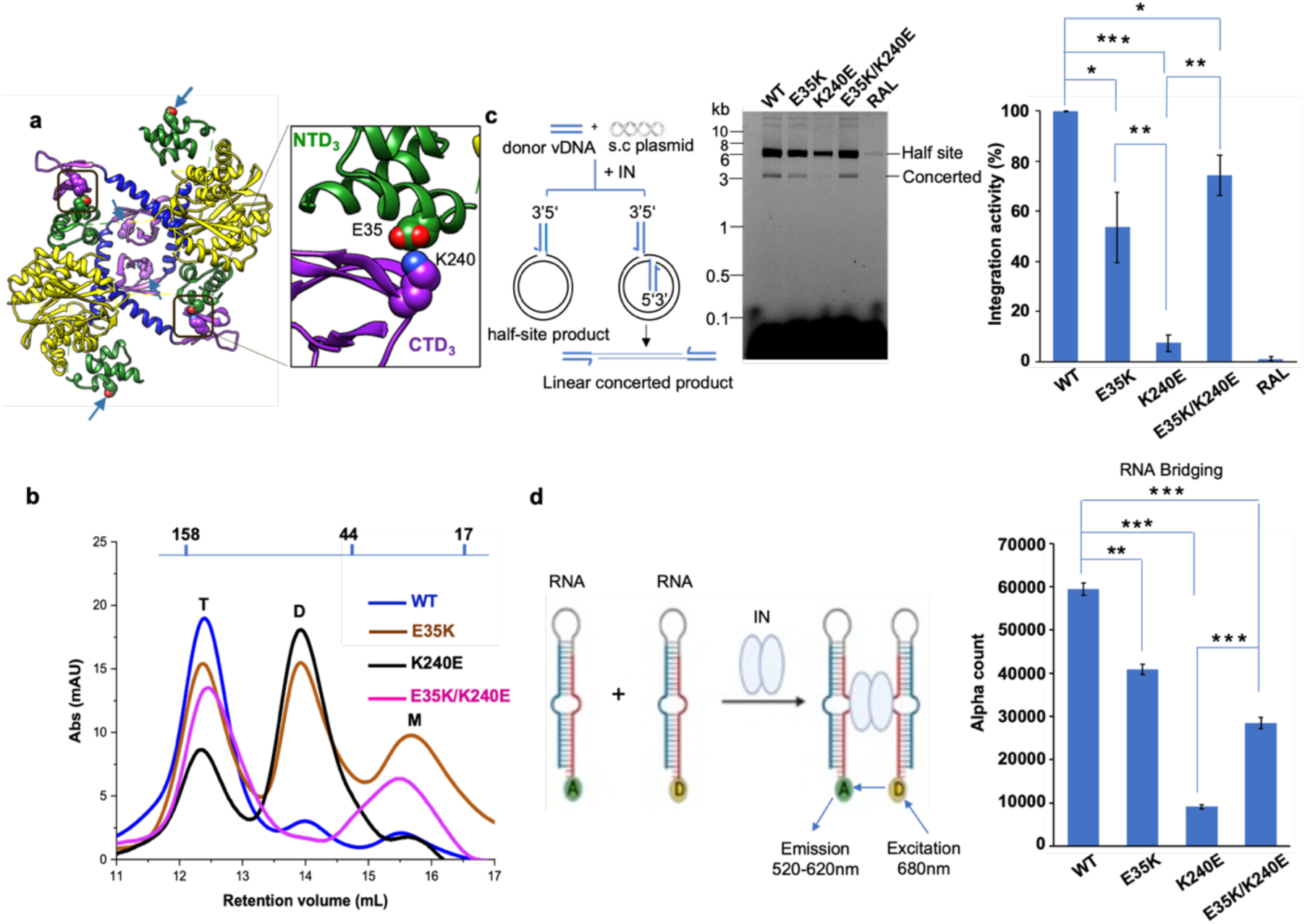
The E35K change restores K240E IN functionality in the context of charge-swapped E35K/K240E *in vitro*. (**a**) Salt bridge interactions formed between E35 in NTD_1_/NTD_3_ and K240 in CTD_1_/CTD_3_ within the outer two regions of the IN tetramer (the inset shows details of the NTD_3_-CTD_3_ interaction). Blue arrows point to the locations of the two other sets of E35 and K240 residues that do not interact with one another and are surface-exposed. (**b**) SEC analysis demonstrating the restoration of K240E tetramerization by the E35K change within the context of the E35K/K240E mutant (T: tetramer, D: dimer, M: monomer). (**c**) Concerted integration assay. Left panel: schematic representation of the assay using vDNA ends, supercoiled (s.c) plasmid DNA, and purified IN. Both half site and concerted integration products are generated as outputs. Middle panel: integration products were detected on 1.5% agarose gel using a fluorescence scanner. The bands corresponding to half site and concerted products are indicated. Right panel: bar graph showing the quantification of concerted integration within the middle panel. RAL, WT IN reactions in the presence of raltegravir IN inhibitor. The integration activity of WT IN was set to 100%, and the activity of mutant IN is presented as a percentage of the WT. The error bars represent SDs of independent experiments, n = 3, each performed with triplicate samples. (**d**) Left panel: Schematic of an AlphaScreen-based RNA bridging assay^26^. Each oval corresponds to an IN multimer. “A” and “D” indicate anti-digoxin Acceptor and streptavidin coated Donor beads bound to either digoxin (DIG) or biotin, respectively. Right panel: quantification of purified IN protein interactions with HIV-1 TAR RNA. Average values are from three independent experiments, with error bars indicating SDs. Statistical significance was determined by the Student’s *t*-test (* P < 0.05, ** P < 0.01, *** P <0.001).

We designed reverse-charge amino acid substitutions E35K, K240E and the charge-swapped double mutation E35K/K240E, expressed the mutant IN proteins in *E. coli* cells, and purified them using nickel affinity and heparin chromatography (**Supplementary Fig. 18**). The purified proteins were analyzed by SEC to assess their multimeric states. Individual IN changes E35K and K240E led to modestly and dramatically reduced levels of the tetramer, respectively. Strikingly, the E35K change in the context of the charge-swapped double mutant substantially restored K240E IN tetramerization (**Figure 4b**). We next examined the effect of these changes on the ability of IN to catalyze two-end vDNA integration *in vitro*. We found that the catalytic activities of mutant INs tracked closely with their ability to form IN tetramers. The concerted integration products were reduced for E35K IN and markedly impaired for the K240E IN. However, the charge-swap double mutant E35K/K240E was substantially more active than its K240E counterpart (**Figure 4c**). Additionally, we examined the effect of these mutations on the ability of IN to bind and bridge vRNA^26,28^. We used the AlphaScreen-based assay^26^ to probe IN-RNA binding and bridging as a proxy for IN’s role in facilitating the localization of RNPs into capsid lattices. Again, the ability of IN to bind and bridge RNA closely correlated with the tetramerization patterns of the mutant proteins. K240E IN was substantially compromised for bridging RNA molecules, whereas E35K and E35K/K240E IN exhibited partly reduced RNA bridging activity (**Figure 4d**). Collectively, these experiments indicate that the tetrameric organization of IN, maintained in part through interactions between E35K and K240E, is relevant for performing the two distinct functional roles of IN *in vitro*.

We next analyzed the consequences of E35K, K240E, and E35K/K240E IN changes in the context of HIV-1 replication. Mutations were introduced into the IN-coding portion of a single-round luciferase reporter virus. Previously analyzed class I and class II IN mutant viruses D64N/D116N and H12N, respectively, were included for comparison^23,64,65^. Viruses were assessed for particle release from transfected HEK293T cells by p24 ELISA, and p24-normalized levels of WT and mutant viruses were used in subsequent infection assays. Mutant viral production was reduced at most 2-fold compared to WT HIV-1 (**Figure 5a**). As previously reported, the K240E and E35K changes instilled severe (>1,000-fold) and comparatively mild (∼2- fold) infection defects, respectively (**Figure 5b**)^8^. Although the E35K/K240E double mutant was also significantly defective (∼300-fold), we note that the addition of the E35K change boosted K240E infectivity and hence the completion of the early events of the viral replication cycle by ∼6- fold. To assess potential contributions from viral DNA synthesis, late reverse transcription (LRT) products were determined by quantitative PCR (qPCR) at 8 h post-infection. While the class II IN mutant control H12N supported ∼5% of WT LRT, the class I mutant D64N/D116N, as expected, supported the WT level of DNA synthesis^24^; E35K also synthesized DNA at the WT level. The K240E mutant, akin to H12N, supported low LRT levels (∼6% of WT). The addition of the E35K mutation increased the efficiency of K240E reverse transcription by ∼3-fold (**Figure 5c**).

**Figure 5.**
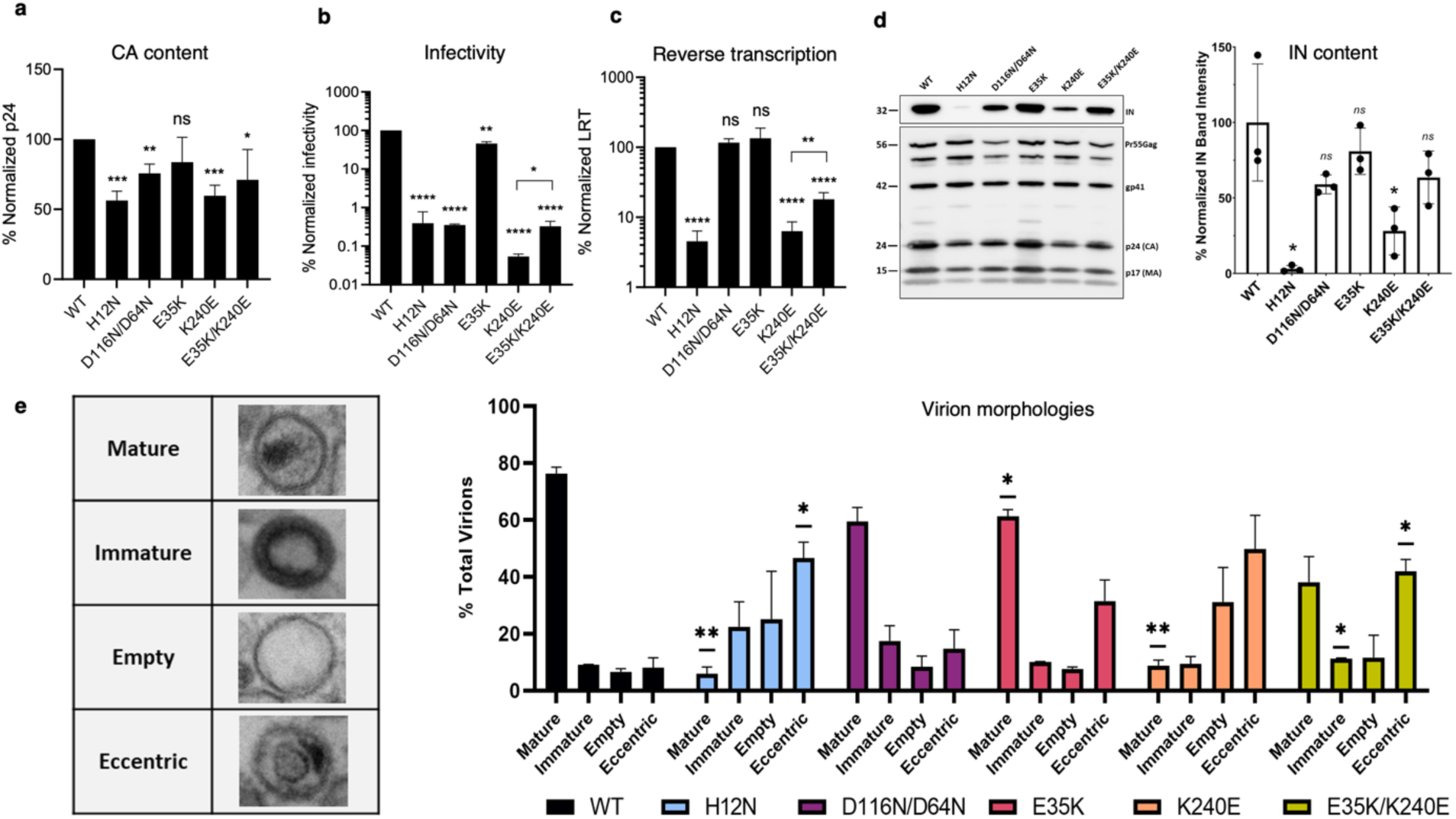
**E35K partially restores K240E IN defects during early and late stages of HIV-1 replication.** (**a**) IN mutant p24 values normalized to WT. (**b**) IN mutant infectivities (RLU/μg) normalized to WT. (**c**) LRT values normalized to WT. (**d**) Immunoblot analysis of IN and Gag/p24 processing intermediates. Representative anti-IN and anti-CA blots from three independent experiments are shown; numbers to the left are mass standards in kDa. Right panel, quantification of IN mutant band intensities relative to p24/p25. (**e**) Frequencies of WT and IN mutant virus morphologies. For each experiment, >200 virions were counted. Categories were assigned based on the following criteria (1) Mature; vRNPs congruent with the CA lattice, (2) Immature; radially arranged electron density, (3) Empty; membrane with no contents, (4) Eccentric; vRNPs outside the CA lattice. Results are: avg. ± SD of n=2 experiments (panels a-c); avg. ± SEM of n=3 experiments (panel d); avg. ± SEM of n=2 experiments (panel e). Statistically significant differences in comparison to WT virus or indicated mutant pairs are represented with asterisks (panels a-d, two-tailed t test; panel e, unpaired t-test followed by multiple comparisons correction; ns, P > 0.05; *, P < 0.05; **, P < 0.01; ***, P < 0.001; ****, P < 0.0001).

To assess effects of the IN changes on HIV-1 structure, we first quantified IN protein content of virion extracts by immunoblotting. As described previously^23,28^, H12N IN was unstable in virions. While the ∼4-fold reduction observed in K240E virions was also statistically significant (**Figure 5d**), the IN content of the remaining viruses was generally similar to WT. Thus, the addition of the E35K change increased the steady-state level of K240E virus IN by about 2-fold. The RNP complex presents as electron-density by transmission electron microscopy (TEM), and ALLINI treatments and class II IN mutations yield predominantly eccentric virions with the RNP situated outside of comparatively lucent capsid structures^26,28,37,66^. As expected, the vast majority of WT particles harbored centrally located electron density congruent with conical cores (**Figure 5e**, left; “Mature”); we also noted comparatively minor populations of immature, empty, and eccentric particles within the WT viral preparations. Whereas class I D64N/D116N and E35K particles appeared broadly similar to the WT, H12N revealed the sharp increase in eccentric particles typical of class II mutant virions. As K240E virion morphologies largely mimicked H12N, we conclude that K240E is a replication-defective class II IN mutant virus. The addition of the E35K change significantly improved the proportion of mature K240E mutant particles. We conclude the charge swapped mutation partially corrects K240E virion structure by increasing the steady state level of IN protein and proportion of mature virion particles.

### Implications for vRNA binding by the IN tetramer

The arrangement of the tetrameric IN assembly has important implications for a working model of its binding to RNA. The four CTDs, which are crucial for RNA binding^26–28^, zigzag along the interface of two IN dimers (**Figure 6a**). These CTDs provide a highly positively charged surface stretching over 80 Å, which could effectively bind RNA (**Figure 6b**). The central cavity (∼40 Å) comprised of inner CTD_2_ and CTD_4_ is suitably sized to engage both single-stranded and double stranded RNAs, while CTD_1_-CTD_4_ and CTD_3_-CTD_2_ are connected through narrow channels that could only accommodate single-stranded segments. Distinct vRNA structural features, including stem/bulges relevant to IN binding^26^, would be expected to dictate engagement of these sites on IN. The structure of the IN tetramer reveals a putative vRNA binding channel and provides initial clues for experimental observations that IN tetramers, rather than dimers (where two CTDs are completely separated from one another) or monomers, can effectively bind vRNA *in vitro* and in virions^26,28^. Class II IN mutant proteins exhibit a pronounced defect in IN’s ability to bind and bridge RNA molecules, in part because they compromise IN tetramerization.

**Figure 6.**
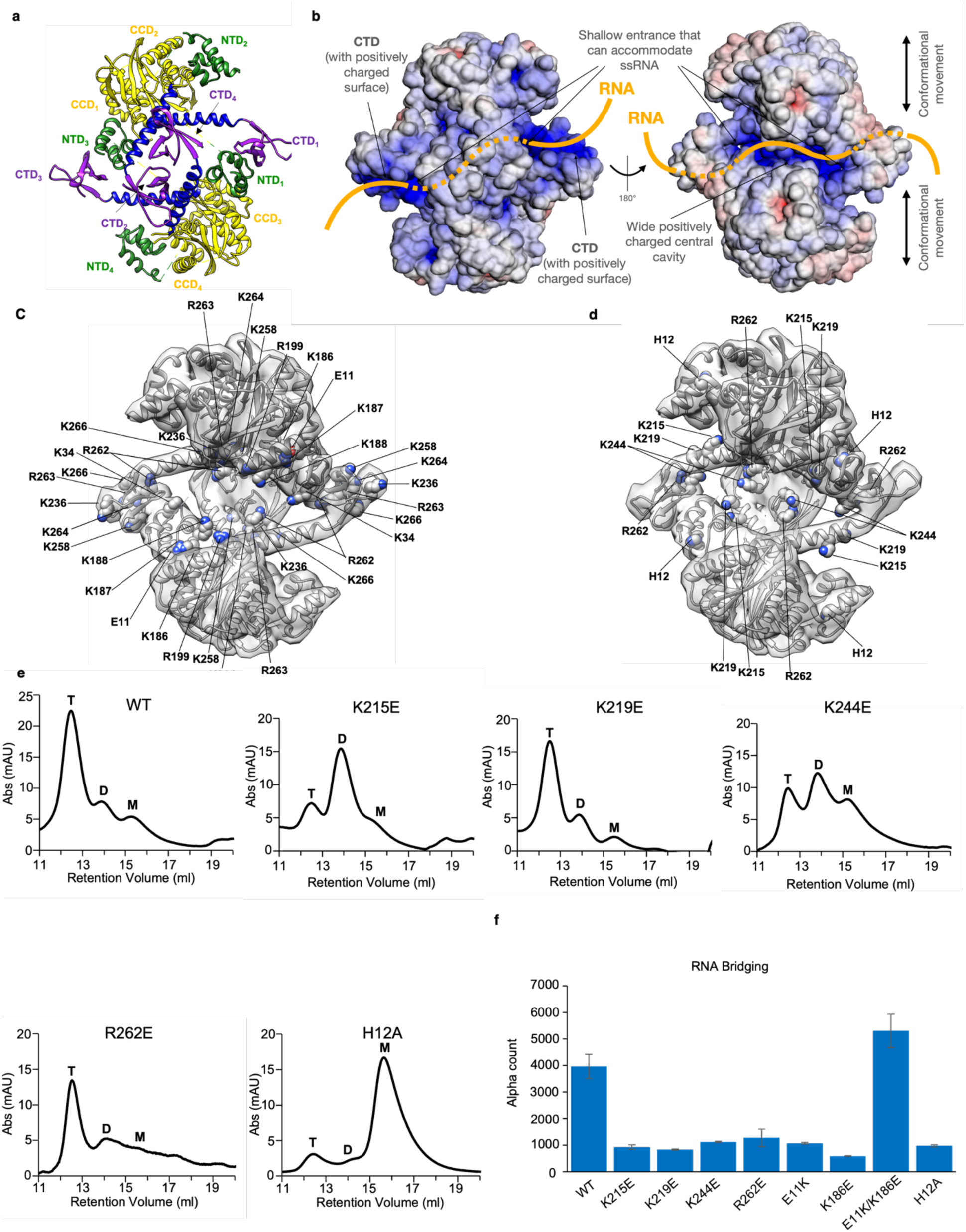
The putative RNA-binding cleft within the IN tetramer. (**a**) Atomic model of the IN tetramer, colored by domain. The four CTDs, which are crucial for RNA binding^26,28,29^, zigzag along the interface of two IN dimers. (**b**) The model is colored by the electrostatic charge distribution. The organization of the CTDs provide a highly positively charged surface stretching over 80 Å, which could effectively bind RNA. A ∼40 Å central cavity comprised of inner CTD_2_ and CTD_4_ is suitably sized to engage both single- and double-stranded RNAs. The putative path of RNA binding to the IN tetramer is shown in orange. **(c)** IN mutations previously analyzed in the literature^26–28^ mapped to the positively charged cleft within the IN tetramer. **(d)** IN mutations newly tested in this work mapped to the positively charged cleft within the IN tetramer. **(e)** SEC profile of WT and mutant IN proteins, showing their multimerization state (T: tetramer, D: dimer, M: monomer). **(f)** Quantification of RNA bridging mediated by purified IN proteins using the AlphaScreen-based RNA bridging assay^26^. Average values are from two independent experiments, with error bars indicating SDs.

We next sought to test whether the positively charged cleft within tetrameric IN affects RNA bridging. Since RNA is negatively charged, we reasoned that mutating positively charged Arg/His/Lys residues to Glu in the vicinity of the cleft would abrogate RNA bridging either by (1) compromising IN tetramerization or (2) directly interfering with charged vRNA/IN interactions. Similar experiments have been previously performed, but those experiments were not guided by the native structure of tetrameric IN^26–28^. **Figure 6c** depicts the location of residues targeted by prior mutagenesis experiments in the context of the positively charged cleft. In all cases, mutations of the positively charged residues compromised RNA bridging^26–28^. To substantiate prior findings, we generated Glu mutants for the remainder of the positively charged residues in the vicinity of the cleft, namely K215E, K219E, K244E, and R262E, depicted in **Figure 6d**, and purified the corresponding IN proteins. We also purified the well-known IN mutant H12A, which is defective for tetramerization, as well as mutants E11K, K186E, and the charge swapped E11K/K186E double mutant. Like the charge-swapped E35K/K240E IN double mutant, the E11K/K186E IN double mutant was previously shown to restore IN tetramerization in comparison to either E11K or K186E alone^18^, although its effect on RNA bridging remained unknown. As expected, H12A was fully defective for tetramerization (**Figure 6e**). While IN mutants K215E and K244E were partially compromised, K219E and R262E changes had little to no effect on IN tetramerization. Notably, all mutant proteins were defective for RNA bridging activity (**Figure 6f**). The charge-swapped E11K/K186E changes restored both IN tetramerization^18^ and RNA bridging activity (**Figure 6f**), as we observed for E35K/K240E, substantiating the evidence that tetramerization is necessary to bind vRNA. As previously suggested^28^, the results point to two possible mechanisms by which IN mutations may compromise RNA binding *in vitro*, either by: (1) affecting functional IN tetramerization or (2) directly compromising IN-RNA interactions. In the cell, mutations may also reduce the level of virion IN, precluding formation of RNPs.

Taken together, our findings strongly support the idea that proper IN tetramerization is an essential prerequisite for IN-vRNA binding in virions and plays a crucial role in IN’s function during virion morphogenesis. The IN tetramer structure provides a plausible model for the RNA-binding cleft and should also help to guide efforts to optimize clinically-relevant allosteric inhibitors^29^.

## Discussion

### HIV-1 IN structural dynamics help choreograph complex functions

HIV-1 IN is a highly dynamic protein^17,18^. Purified IN protein stoichiometry generally ranges from monomer to tetramer, with the tetramer being favored at higher protein concentrations or at low salt. Although the native form of HIV-1 IN in virions is likely tetrameric,^29^ the structure of this inherent IN multimer has resisted resolution despite three decades of IN structural biology efforts. Here, we solved the structure of the IN tetramer stabilized by the LEDGF/p75 IBD and elucidated snapshots for how the tetramer rearranges to assemble the intasome hexadecamer. Our independent structure of the apo IN tetramer lacking the IBD indicated that the host factor fragment did not significantly impact global IN organization within the functional multimer. Whereas the conformational changes necessary for intasome formation within and between individual IN domains, mediated by flexible NTD-CCD and CCD-CTD linkers, have been well-described and are generally conserved^6–8,10,59–62^, changes to the tetrameric organization of lentiviral INs have remained unknown. Our structural biology data implies that the two IN dimers must contract while tethered through the CTD scaffold. An analogy would be the wing movement of a butterfly, with the tetrameric CTD arrangement representing the body and the two NTD:CCD dimers the wings. IN dynamics about the central interlocked CTDs, which are expected to be more pronounced in the absence of the IBD, may also be relevant to engaging/disengaging the viral RNA and for recognizing diverse RNA lengths. The data also informs a better understanding for why intasomes from HIV-1 (and likely by extension SIV) are more pleiotropic than those from MVV. Both hexadecameric intasome assemblies maintain bridging CTD:CTD interactions between tetramers I/III and II/IV, respectively; however, the inter-tetramer interaction interface made by the CTDs is weaker in HIV-1 in comparison to MVV (**Supplementary Fig. 17**). Thus, it is not surprising that the flanking HIV-1 INs frequently dissociate *in vitro*, in turn making the whole complex compositionally pleiotropic. Collectively, these data add to our understanding of IN structural dynamics and suggest mechanisms by which these properties may help choreograph its complex functions during HIV-1 replication.

### The oligomeric forms of HIV-1 IN relevant to function

Since the discovery of IN, there have been ongoing debates about its oligomeric form required for catalytic integration. Structures of PFV intasomes contained four IN protomers assembled around vDNA ends^6^. However, the first structures of HIV-1 intasomes complicated interpretations; whereas the conserved tetrameric core was analogous to PFV, larger species, including octamers, decamers, and dodecamers, were identified in particle subsets by cryo-EM^8^. The intasome from MVV, which is biochemically more stable and hence more homogeneous than the HIV-1 complex, was a hexadecamer, although smaller species were also observed^10^. The pleiotropic nature of lentiviral IN / intasomes is well-established^8,10,12,19^. Loss-of-function experiments using MVV IN and corresponding viruses showed reduced ability of IN mutants to assemble into tetramers and form intasomes to catalyze strand transfer, with concomitant defects in viral infectivity^9^. Analogously, numerous loss-of-function and mutagenesis studies suggested that the composition of IN required for RNA binding is tetrameric. However, all loss-of-function experiments are inherently inconclusive because mutations may have pleiotropic effects, thereby complicating conclusions. Thus, both the nature of the multimeric lentiviral IN assembly relevant for integration and for vRNA binding have remained unclear. In this regard, our identification of a key salt-bridge interaction between residues E35 and K240, and *in vitro* experiments with E35K, K240E and E35K/K240E mutant INs demonstrating that both catalytic activity and RNA binding are rescued by the double mutant and closely correlate with tetramerization, are noteworthy. In regard to catalytic activity, the simplest explanation is that compromising tetramerization would predominantly affect intasome assembly, and indirectly affect function, since it was shown that, once assembled, different intasome forms are equally active for concerted integration *in vitro*^15^. Additionally, our virology experiments with these mutants highlight the indispensable role of the IN tetramer for viral infectivity. We note that the effects of E35K and K240E mutations alone are non-equivalent, and the phenotypic effect of K240E is more severe across all experimental assays. Although the introduction of either individual E35K or K240E mutations is expected to comparably compromise the salt-bridge interaction, there are two additional sets of E35/K240 residues in the tetramer structure outside the salt-bridge interface (**Figure 4a**). The effect of K240E likely therefore extends beyond the salt bridge interface to exert pleiotropic effects on both intasome formation and on IN-mediated RNA binding, which would affect HIV-1 particle morphogenesis. HIV-1 intasomes are biochemically more pleomorphic than corresponding MVV complexes. Although such pleomorphic biochemical complexes are similarly active for concerted integration in vitro, we believe the hexadecamer, based on analogy to other lentiviral intasomes such as the well-characterized hexadecameric MVV intasome (**Supplementary Fig. 17**), is the likely higher-order form relevant in the context of HIV-1 infection. Collectively, we conclude that in the context of cellular infection, the most likely higher-order form of the HIV-1 intasome is hexadecameric, and the relevant form of HIV-1 IN for vRNA binding is tetrameric.

### Study limitations

Because our structures were based on IN oligomers assembled from purified, recombinant components and not complexes isolated from infected cells, we can only infer biological relevance. Critically, our mutagenesis strategy indicates that the E35K-K240E salt bridge and, by extension, the IN tetramer structure, is biologically relevant. At the same time, our ability to inform the pathway of intasome assembly from the tetramer structure is limited through comparison of two independently resolved cryo-EM structures. Similarly, although our data strongly support the conclusion that intasomes that fail to multimerize will exhibit compromised integration efficiency, we cannot unequivocally know the oligomeric form of intasomes in cells, especially under conditions of stress. For example, a prior report has suggested that there may be dynamic changes between the number of INs within pre-integration complexes that is dependent on their cellular location, interaction with LEDGF/p75, and/or exposure to ALLINIs^67^. Although our model represents the most straightforward and accordingly likely biologically relevant pathway, it does not consider more elaborate scenarios that, for example, might require tetramer disassembly into composite components to assemble an active HIV-1 intasome.

## Materials and Methods

### Protein expression and purification of WT IN and IBD complex

The pCPH6P-IN plasmid contains full-length of HIV-1 NL4-3 IN with a N-terminal 6His tag, followed by a HRV 3C protease cleavage site. The pES-IBD-3C7 plasmid contains the LEDGF/p75 IBD with a N-terminal 6His-SUMO tag, followed by a SUMO protease cleavage site. These two plasmids were co-transformed into *E. coli* Rosetta DE3 competent cells (NEB) to co-express full-length HIV-1 IN and the IBD. The cells were cultured in LB medium supplemented with 50 µM ZnCl_2_ and appropriate amount of antibiotics kanamycin and ampicillin. When the OD of the culture reached ∼1.0 at 37°C, the temperature was dropped to 16°C, and expression was induced overnight with the addition of 0.4 mM isopropyl β-D-1-thiogalactopyranoside (IPTG). Following overnight expression, cells were harvested for protein purification by centrifugation at 6,000 *g*.

To purify the IN-IBD complex, the cells were sonicated in a buffer containing 1 M NaCl, 20 mM imidazole, 0.2 mM PMSF, 50 mM Tris-HCl, 5 mM 2-mercaptoethanol, pH 7.6, then the cell lysate was subjected to centrifugation at 39,000 *g* for 30 minutes. The supernatant was immediately loaded into a 5 mL HisTrap column (Cytiva), which was attached to a FPLC system for purification. The complex was washed at 8% and 20% buffer B containing 250 mM imidazole to remove non-specific bound proteins, followed by an imidazole gradient wash and elution from 20%-100% of buffer B (1 M NaCl, 250 mM imidazole, 50 mM Tris-HCl, pH 7.6). The 6His tag on IN and the 6His-SUMO tag on the IBD were cleaved in the presence of 5 mM 2-mercaptoethanol using SUMO and HRV 3C proteases, respectively. The protein solution was then subjected to another 5 mL HisTrap column purification to remove the cleaved tags. The final IN/IBD complex was purified by SEC using a Superdex-200 column in 750 mM NaCl, 50 mM Tris-HCl, pH 7.6, 5 mM DTT and 10% glycerol.

In general, we found that the IN-IBD complex does not need additional buffer components, such as CHAPS detergent, to remain soluble and tetrameric during purification. In contrast, when we purified IN alone, all the initial steps, including resuspension, lysis, and HisTrap and heparin column purifications, required CHAPS to be present.

### Cryo-EM sample preparation and data collection for the IN-IBD tetramer

IN-IBD complex (0.1 mg/mL) was applied onto freshly plasma cleaned (20s, Solarus plasma cleaner) holey grids (Quantifoil UltrAufoil R 1.2/1.3 300 mesh) at 4°C in the cold room with ∼90% humidity. The sample was settled onto grids for 30 sec before manually blotting for 8 sec, then plunged into the liquid ethane using a manual plunger. The vitrified grids were transferred to liquid nitrogen for storage and data collection.

The IN-IBD complex cryo-EM dataset was collected on a Talos Arctica transmission electron microscope (Thermo Fisher Scientific) operating at 200 keV, equipped with K2 direct electron detector (Gatan) with a GIF Quantum Filter with a slit width of 20 eV. The data collection was performed using Leginon^68,69^ with the stage tilt set to 30**°**–50°^46,48,70^. Movies were recorded at a nominal magnification of 45 kx, corresponding to a calibrated pixel size of 0.92 Å. The total dose for this dataset is 26.5 e^-^/ Å^2^, using dose rate of 22.7 e^-^/ pix/s. The imaging parameters for this dataset are summarized in **Supplementary Table 1**.

### Graphene grid preparation for the apo IN tetramer cryo-EM dataset

UltrAufoil R1.2/1.3 grids (Quantifoil) were plasma cleaned using the Solarus II plasma cleaner (Gatan) with an argon-oxygen recipe as follows: Ar flow = 35 sccm; O_2_ flow = 11.5 sccm; power = 40W, time = 30 s. Whatman filter paper supported on a wire mesh was submerged in a trough filled with Milli-Q water, to which plasma cleaned grids were then gently transferred with the gold foil facing top side. We then transferred 1cm x 1cm Trivial Transfer Graphene (ACS) onto the trough containing water and the grids, based on manufacturer’s instructions. The graphene supported on the sacrificial layer of poly(methyl methacrylate) (PMMA) was gently layered along onto the grids and then the grids were carefully transferred onto another Whatman filter paper for quick drying. After air drying, the grids were incubated at 300°C for 60 min. Grids containing a thin layer of graphene and PMMA were transferred onto a crystallizing dish with a Pelco TEM Grid Holde holder submerged in acetone. Grids were washed thrice in acetone to dissolve PMMA with washes ranging from 1 h to overnight with gentle shaking (<20 rpm). A final wash was performed with fresh isopropanol for 30 min with gentle shaking. After discarding the isopropanol, the grids were dried on Whatman filter paper in a fume hood, then transferred to a petri dish with Whatman filter paper and dried in an oven at 200°C for 20 min. After drying, grids were immediately transferred to grid boxes and vacuum sealed until they were used for specimen preparation. Prior to sample application, grids were plasma cleaned using a hydrogen-oxygen recipe as follows: H flow = 30 sccm; O_2_ flow = 11.5 sccm; power = 10W, time = 10 s.

### Cryo-EM sample preparation and data collection for the apo IN tetramer

Apo IN (0.04 mg/mL) was applied onto freshly plasma cleaned (20s, Solarus plasma cleaner) gold grids (Quantifoil UltrAufoil R 1.2/1.3 300 mesh) coated with a thin layer of graphene at 4°C in the cold room with ∼90% humidity. The sample was settled onto grids for 1 min prior to manually blotting for 2 sec, then plunged into the liquid ethane using a manual plunger. The vitrified grids were transferred to liquid nitrogen for storage and data collection.

The apo IN cryo-EM dataset was collected on a Talos Arctica transmission electron microscope (Thermo Fisher Scientific) operating at 200 KeV, equipped with Falcon 4i direct electron detector. The data collection was performed using EPU. Movies were recorded at a nominal magnification of 105kx, corresponding to a calibrated pixel size of 0.95 Å. The total dose for this dataset is 60 e^-^/ Å^2^ using dose rate of 10 e^-^/pix/s. The imaging parameters for the dataset are summarized in **Supplementary Table 1**.

### Cryo-EM image processing for the IN-IBD tetramer and apo IN tetramer

All data processing for both the IN-IBD tetramer and the apo IN tetramer were performed in cryoSPARC V4.7.0^49^ . For the IN-IBD dataset, the movies were preprocessed using patch-based motion correction with a gain reference that was generated during data collection, followed by patch-based CTF estimation. Micrographs for which CTF estimates were poorer than 10 Å were discarded. The remaining micrographs were used for particle selection with the blob picker modality in cryoSPARC. The selected particles were manually inspected, and bad particle selections were removed by adjusting the threshold of NCC score and local power. The remaining particles were extracted from micrographs using a box size of 300 pixels. 2D classification was then performed using 40 iterations and a particle batch size of 200, which helps with extracting signal for small particles characterized by low signal-to-noise ratios. The 2D class averages that produced protein features were used for template picking with a particle diameter of 150 Å. The selected particles were inspected and extracted from micrographs. The extracted particles were subjected to iterative 2D classification to remove bad particles that did not produce class averages containing protein features. The resulting particles from the last round of 2D classification were subjected to *ab initio* reconstruction with 2 classes and C2 symmetry applied. One of the *ab initio* reconstructions that showed distinct features consistent with the IN-IBD structure was selected for non-uniform refinement^71^, with C2 symmetry applied. The resulting dataset then underwent reference-based motion correction followed by global CTF refinement. We also attempted to perform local CTF refinement, but the map quality and the nominal resolution did not improve. Therefore, local CTF refinement was not employed for the generation of the final map. The map produced after reference-based motion correction and global CTF refinement was input into one last round of non-uniform refinement, which generated a 4.1 Å map used for model building (**Supplementary Table 1** and **Figure 1**). The complete workflow is shown in **Supplementary** Figure 3, and the validation of the map is shown in **Supplementary** Figure 4.

The data processing steps for the apo IN tetramer were similar to those employed for IN- IBD, with the following exceptions: a model of ResNet16 within Topaz^72^ was retrained using a minibatch size of 256 with selected 2D class averages from an initial round of classification, and the Topaz extract modality with this retrained model was employed for particle selection. Following iterative 2D classification, *ab initio* reconstructions with 3 classes and C2 symmetry were performed. The particles from two of the three *ab initio* maps were combined and subjected to heterogeneous refinement with 3 classes and C2 symmetry applied. Reconstructions were judged based on the Fourier shell correlation (FSC) criteria, as well as the helical density for the CCD-CTD linkers; if the linkers were broken, this indicated that the complex was unstable, but if the linkers were intact, this indicated that the complex maintained its structural integrity. The particles and the map corresponding to the best reconstruction were used as input for non-uniform refinement, followed by global CTF refinement and another round of non-uniform refinement. These steps produced a 7 Å map (**Supplementary Table 1**). The complete workflow is shown in **Supplementary** Figure 6, and the validation of the map is shown in **Supplementary** Figure 7. A comparison of the apo IN tetramer with the IN-IBD tetramer is displayed in **Supplementary** Figure 8.

The directional resolution of both maps was initially evaluated using the 3D FSC server (3dfsc.salk.edu)^46^, and the quality of the orientation distribution was evaluated using the Sampling Compensation Function (SCF)^73,74^. The validation images displayed in the supplementary figures were obtained directly from cryoSPARC. All images were made using UCSF Chimera^75^.

### Model building for the IN-IBD tetramer

The atomic model of HIV-1 IN tetramer was constructed in a stepwise manner. We first docked all individual domains into density, beginning with high-resolution crystal structures of the CCD:CCD dimer^51^, the NTD^76^, and the CTD^57^, which were rigid body docked into the 4.1 Å cryo-EM map using UCSF Chimera. The alpha helical linkers connecting the CCD to the CTD were built automatically using Rosetta^77^. Subsequently, the model was subjected to one round of real-space refinement in the Phenix suite using phenix.real_space_refine with restraint weights^78^. The model was manually adjusted in Coot^79^, and the discrepant parts in the model were removed. Phenix and Coot were used iteratively during model building and refinement. The model was refined through one last round of real-space refinement in the Phenix suite to generate the final model. The statistics of the final model are shown in **Supplementary Table 1**.

### Crosslinked HIV-1 Intasome preparation

Intasomes containing pre-cleaved vDNA ends were assembled using essentially the same conditions as for the concerted integration assay descried above, except the target plasmid DNA was omitted, 50 μM dolutegravir was present in the reaction buffer, and the vDNA substrate was substituted by U5-69T, which was prepared by annealing LTR-29 (5’- AAAAAAAAGTGTGGAAAATCTCTAGCA-3’) with U5-69R (5’- ACTGCTAGAGATTTTCCACACTTTTTTTTTTTTTTTTTTTTTTTTTTTTTTTTTTTTTTTTTTTTTTTTTTTTTTTTTTTTTTTTTTTTT-3’). The reaction was carried at 37°C for 120 min and stopped on ice, then, NaCl (1.0 M final) was added to the reaction mixtures to solubilize the intasomes. Purification of intasomes was essentially as previously described^13^. The reaction mix was loaded onto a HisTrap column (GE Healthcare) equilibrated with 20 mM HEPES pH 8.0, 5 mM 2- mercaptoethanol, 1.0 M NaCl, 20% (w/v) glycerol. Intasomes were then eluted with a linear gradient of 0 mM to 500 mM imidazole in the same buffer. Pooled intasomes were crosslinked by 2.0 mM BS3 (Pierce). The crosslinking reaction was quenched by adding 100 mM Tris, pH 8.0. Aggregates/stacks of intasomes and free IN were then removed by gel filtration on a TSKgel UltraSW Aggregate HPLC column (Tosoh Bioscience) in 20 mM Tris pH 6.2, 0.5 mM TCEP, 1.0 M NaCl, 5.0 mM MgCl_2_, and 10% (w/v) glycerol. Intasomes corresponding to single species were concentrated to 0.4 mg/ml for EM studies.

### Cryo-EM sample preparation and data collection for the HIV-1 intasome

Intasome complex (0.4 mg/mL) was applied onto freshly plasma cleaned (20s, Solarus plasma cleaner) holey grids (Quantifoil UltrAufoil R 1.2/1.3 300 mesh) at 4°C in the cold room with ∼90% humidity. The sample was settled onto grids for 30 sec before manually blotting for 8 sec, then plunged into the liquid ethane using a manual plunger. The vitrified grids were transferred to liquid nitrogen for storage and data collection.

The intasome complex cryo-EM dataset was collected on a Titan Krios transmission electron microscope (Thermo Fisher Scientific) operating at 300 keV. The data collection was performed using Leginon^68,69^ with the stage tilt set to 30**°**^46,48,70^. 775 movies were recorded at a nominal magnification of 29 kx, corresponding to a calibrated pixel size of 1.015 Å. The total dose for this dataset was 60.0 e^-^/ Å^2^, using dose rate of 5.2 e^-^/ pix/s. The imaging parameters for this dataset are summarized in **Supplementary Table 1**.

#### Computational classification and refinement of the hexadecameric HIV-1 intasome

A consensus map of the hexadecameric intasome was generated using the following procedures, performed in cryoSPARC V4.2.1^49^. The movies of crosslinked intasomes (**Supplementary Fig. 13**a) were preprocessed using patch-based motion correction with a gain reference that was generated during data collection, followed by patch-based CTF estimation. Micrographs for which CTF estimates were poorer than 10 Å were discarded. The remaining micrographs were used for particle selection with the blob picker modality in cryoSPARC. The selected particles were manually inspected and bad particle selections were removed by adjusting the threshold of NCC score and local power. The remaining 238,303 particles were extracted from micrographs using a box size of 320 pixels. 2D classification was performed using default parameters. The 2D class averages that produced intasome features were selected (**Supplementary Fig. 13**b), corresponding to 147,569 particles. These particles were subjected to *ab initio* reconstruction with 2 classes and C2 symmetry applied. One of the *ab initio* reconstructions showed distinct features consistent with the intasome. This map was used for homogeneous refinement with C2 symmetry applied, yielding a reconstruction that resembled the intasome devoid of flanking subunits and resolved to a nominal resolution of 4.0 Å (**Supplementary Fig. 13**c-f and **Supplementary Table 1**). This resulting map was used as the input for a multi-body refinement procedure that we implemented within cryoSPARC.

Based on the assumption that the two flanking regions are independently flexible, we proceeded with resolving each flanking region within the intasome core separately in cryoSPARC^49^. The map was divided into a “top” half and a “bottom” half, as shown in **Supplementary Fig. 14**. To resolve the top flanking region withing core, 147,569 particles were subjected to density subtraction to first remove signal in the bottom flanking region. The subtracted particles were classified using global 3D classification into five classes. From these classes, 70,680 particles from the best three classes had density within the top flanking region. These particles were selected and subjected to a second round of 3D classification using a mask focused on the top flanking region. From focused classification, we selected 11,658 particles that contained clearly resolved density for the entirety of the flanking region, whereby linker density and density for all domains was evident. These particles were then subjected to local CTF refinement, which was followed by a round of local refinement using a mask on the entirety of the volume. This procedure yielded a final map of the top half of the HIV-1 intasome. We resolved the bottom half of the intasome using an identical method, with 11,122 particles contributing to the final map. The procedure is demonstrated in **Supplementary Fig. 14**. Both maps were validated using global and local resolution estimates in cryoSPARC. The 3DFSC was generated using the 3D FSC server following standard procedures^46,80^. The surface sampling plot for the Euler angle distribution was generated using standalone scripts, as previously described^70,74^. The composite map was stitched together from two independently refined halves of the intasome after aligning each half to the intasome core and summing the two maps using maximum pixel intensities as the summation criterion. The validation for both maps and the generation of the composite map is shown in **Supplementary Fig. 15**. The visualization of all maps was performed in UCSF Chimera and ChimeraX^75,81^.

### Construction of the rigid bhody docked model for the hexadecameric HIV-1 intasome

We used a rigid body docking protocol to construct the hexadecameric model of the HIV- 1 intasome. The model was constructed in a stepwise manner. In some instances, multiple different orientations of a domain could be docked into density, or the docked domain was not constrained within the region of interest in the density. This occasionally was observed when docking individual CTDs. In these instances, we segmented the experimental cryo-EM density to include only the region of interest and additionally used a portion of the extended CCD-CTD linker to better orient and register the model within the segmented density. The methods here describe one half of the intasome complex, corresponding to protein chains A through H, and DNA chains Q-R. First, the structure of the cleaved synaptic complex intasome (PDB 6PUT) was docked into the core, yielding chains A-B of the inner IN dimer and the CTDs of chains C and E. The outer IN dimer was constructed by docking the structure of the two-domain HIV-1 IN NTD-CCD (PDB 1K6Y) and the two-domain IN CCD-CTD (PDB 1EX4) into the density. Connectivity of the NTDs and the CTDs was verified by inspecting the density and comparing with the high-resolution structure of the MVV hexadecameric intasome (PDB 7Z1Z). Both sets of flanking IN dimers were constructed in the same manner, by docking the structure of the two-domain HIV-1 IN NTD-CCD (PDB 1K6Y) and the two-domain IN CCD-CTD (PDB 1EX4) into the density, then verifying the connectivity using the structure of the MVV intasome (PDB 7Z1Z). Additionally, the interaction between the CTDs of chains F and H were confirmed using the structure of the CTD:CTD dimer (PDB 1IHV), and the interaction between the NTD and CTD of chain G was confirmed using the structure of the cleaved synaptic complex intasome (PDB 6PUT, chain B), whereby the same interfaces are engaged. Docking for chains G and H, and the CTD of chain F, which correspond to the lowest resolution regions of the map, are also shown in **Supplementary Fig. 16**a. A table depicting the PDB IDs that were used in the procedure is shown below:

|  | NTD (PDB codes) | CCD (PDB codes) | CTD (PDB codes) |
| --- | --- | --- | --- |
| Chain A (tetramer 1) | 6PUT | 6PUT | 6PUT |
| Chain B (tetramer 1) | 6PUT | 6PUT | 6PUT |
| Chain C (tetramer 1) | 1K6Y, 7Z1Z | 1K6Y | 6PUT |
| Chain D (tetramer 1) | 1K6Y, 7Z1Z | 1K6Y | 1EX4, 7Z1Z |
| Chain E (tetramer 2) | 1K6Y, 7Z1Z | 1K6Y | 6PUT |
| Chain F (tetramer 2) | 1K6Y, 7Z1Z | 1K6Y | 1EX4, 7Z1Z, 1IHV |
| Chain G (tetramer 2) | 1K6Y, 7Z1Z, 6PUT | 1K6Y | 1EX4, 7Z1Z |
| Chain H (tetramer 2) | 1K6Y, 7Z1Z | 1K6Y | 1EX4, 7Z1Z, 1IHV |

### Expression and purification of WT and mutant IN proteins

Mutations were introduced into a plasmid backbone expressing 6xHis tagged pNL4-3- derived IN by QuikChange site-directed mutagenesis kit (Agilent). The plasmids containing full-length WT and mutants were transformed into BL21 (DE3) *E. coli* cells. The cells were cultured in LB medium supplemented with 50 µM ZnCl_2_ and appropriate amount of antibiotics kanamycin and ampicillin. When the OD of the culture reached ∼1.0 at 37°C, the temperature was dropped to 16°C, and expression was induced overnight with the addition of 0.4 mM isopropyl β-D-1- thiogalactopyranoside (IPTG). Next day, the cells were harvested by centrifugation at 4200 *g*, 15 min and suspended in Ni buffer A containing 45 mM HEPES, pH 7.5, 6.75 mM CHAPS, 0.9 M NaCl, 10% glycerol, 3 mM 2-mercaptoethanol, 20 mM imidazole. The resuspended cells were sonicated and subjected to centrifugation at 39,000 *g* for 30 min. The supernatant was immediately loaded into a 5 mL HisTrap column, attached to a FPLC system at a flow rate of 1 ml/min. After sample loading, a gradient washing/elution was carried out using buffers A and B (45 mM HEPES, pH 7.5, 6.75 mM CHAPS, 0.9 M NaCl, 10% glycerol, 3 mM 2-mercaptoethanol, 500 mM imidazole). Fractions (1.5 ml) containing target proteins were pooled and diluted to 300 mM salt and loaded onto a Heparin column equilibrated with 30% Heparin buffer B containing 50 mM HEPES, pH 7.5, 7.5 mM CHAPS, 1 M NaCl, 10% glycerol, 3 mM 2-mercaptoethanol a flow rate of 3 ml/min followed by gradient wash from 30% Heparin buffer B to 100%. Fractions across the peak were run on SDS-PAGE and clean fractions containing target proteins were pooled and concentrated for downstream assays.

### In vitro IN concerted integration assay

Integration assays were conducted essentially as described^14^ with slight modifications. Briefly, 2.0 μM IN and 1.0 μM vDNA were preincubated on ice in 20 mM HEPES (pH 7.5), 25% glycerol, 10 mM DTT, 5 mM MgCl_2_, 4 μM ZnCl_2_, 50 mM 3-(Benzyldimethyl-ammonio) propanesulfonate (Sigma), 3.0 μM polyethylenimine (Sigma) and 100 mM NaCl in a 20-μl reaction volume. The vDNA (Integrated DNA Technologies) was fluorescently labeled with a 6- carboxyfluorescein (6-FAM) fluorophore attached to the 5′-end of the oligonucleotide, yielding 5’- Fam-AGCGTGGGCGGGAAAATCTCTAGCA, which was annealed with 5’- ACTGCTAGAGATTTTCCCGCCCACGCT-3’ to form the precut vDNA substrate. Target plasmid DNA pGEM-9zf (300 ng) was then added and the reaction was initiated by transferring to 37°C, and incubated for 2 h. The integration reactions were stopped by addition of SDS and EDTA to 0.2% and 10 mM, respectively, together with 5 μg of proteinase K and further incubated at 37°C for an additional 1 h. The DNA was then recovered by ethanol precipitation and subjected to electrophoresis in a 1.5% agarose gel in 1× tris-boric acid–EDTA buffer. DNA was visualized by fluorescence using a Typhoon 8600 fluorescence scanner (GE Healthcare). Densitometry was used to quantify IN activity. The density of the bands corresponding to concerted integration on fluorescence-scanned gels was quantified by ImageQuant (GE Healthcare). The integration activity of WT IN was set to 100%, and mutant activities are expressed as percentages of WT activity. The error bars represent SDs of independent experiments, n = 3, performed in triplicate. Statistically significant differences in comparison to WT and between experimental groups are represented with asterisks, using the Student’s t-test (* P < 0.05, ** P < 0.01, *** P <0.001).

### Size Exclusion chromatography (SEC)

Purified IN proteins were diluted to 1.5 mg/ml with running buffer containing 20 mM HEPES (pH 7.5), 750 mM NaCl, 10% glycerol, 5 mM CHAPS and 5 mM 2-mercaptoethanol and incubated for 1 h at 4°C followed by centrifugation at 10,000 g for 10 min at 4°C. Multimeric state of supernatant WT and mutant INs were analyzed on Superdex 200 10/300 GL column (GE Healthcare) equilibrated with running buffer at a flow rate of 0.2 ml/min.

For analytical SEC, WT and mutant IN proteins were analyzed using Superdex 200 10/300 GL column (GE Healthcare) with running buffer containing 20 mM HEPES (pH 7.5), 1 M NaCl, 10% glycerol and 5 mM 2-mercaptoethanol at the flow rate of 0.3 mL/min. The protein stocks were diluted to 20 µM with the running buffer and incubated for 1 h at 4 °C followed by centrifugation at 10,000 x g for 10 min. To estimate multimeric state of IN proteins, we used the protein standard including bovine thyroglobulin (670,000 Da), bovine gamma-globulin (158,000 Da), chicken ovalbumin (44,000 Da), horse myoglobin (17,000 Da) and vitamin B12 (1,350 Da). Retention volumes for different oligomeric forms of IN were as follows: tetramer ∼12.5 mL, dimer ∼14 mL, monomer ∼15-16 mL.

### IN-RNA interactions

To monitor IN-RNA interactions we utilized AlphaScreen-based assay^26^, which monitors the ability for IN to bind and bridge between two trans-activation response element (TAR) RNAs. Briefly, equal concentrations (1 nM) of two synthetic TAR RNA oligonucleotides labeled either with biotin or digoxin (DIG) were mixed and then streptavidin donor and anti-DIG acceptor beads at 0.02 mg/mL concentration were supplied in a buffer containing 100 mM NaCl, 1 mM MgCl_2_, 1 mM DTT, 1 mg/mL BSA, and 25 mM Tris (pH 7.4). After 2 h incubation at 4°C, 320 nM IN was added to the reaction mixture and incubated further for 1.5 h at 4°C. AlphaScreen signals were recorded with a PerkinElmer Life Sciences Enspire multimode plate reader.

### Virological assays

To build IN mutant viral constructs, the larger DNA fragment of pNLX.Luc.R-.ΔAvrII^82^ following digestion with PflMI and AgeI restriction enzymes (New England Biolabs) was assembled with 1.85-kb synthetic fragments (Twist Bioscience) containing single or double E35K and K240E changes using NEBuilder HiFi DNA Assembly MasterMix under conditions recommended by the manufacturer. Analogous H12N^65^ and D116N/D64N^64^ mutant constructs were previously described.

HEK293T cells obtained from America Type Culture Collection were grown in Dulbecco’s modified Eagle’s medium containing 10% fetal bovine serum, 100 IU/ml penicillin, and 100 𝜇g/ml streptomycin (DMEM) at 37 °C in the presence of 5% CO_2_. Depending on the assay, viruses were produced at two different scales. For LRT-qPCR and infectivity assays, ∼10^6^ cells were plated in 6-well plates. The following day, cells were co-transfected with ∼2.5 µg total plasmid DNA (pNLX.Luc.R-.ΔAvrII and a vesicular stomatitis virus G envelope expressor at 6:1 ratio using PolyJet™ DNA transfection reagent; SignaGen Laboratories). At 48 h post-transfection, supernatants were centrifuged at 500 x*g* for 5 min, filtered through 0.45 µm filters, and treated with 20 U/ml Turbo DNase for 1 h. The concentration of p24 was determined using a commercial ELISA kit (Advanced Bioscience Laboratories). For immunoblotting and TEM, transfections were scaled up (10^7^ cells plated the previous day in 15 cm dishes) to 30 µg total plasmid DNA. Resulting 0.45 µm filtered supernatants were pelleted by ultracentrifugation using a Beckman SW32-Ti rotor at 26,000 rpm for 2 h.

Cells (∼10^5^ plated in 24-well plates) were infected at 0.25 pg/cell. After 2 h, virus was removed and cells were replenished with fresh DMEM. At 48 h post-infection, cells were lysed with passive lysis buffer (Promega) and luciferase activity, assessed as relative light units per µg of total protein in the cell extracts (RLU/𝜇g), was determined as previously described^83^. For LRT- qPCR, parallel infections included dimethyl sulfoxide vehicle control or 10 µM efavirenz (EFV) to control for potential plasmid carryover from transfection. Genomic DNA was isolated at 8 h post-infection using Quick-DNA™ Miniprep Kit (Zymo Research). Quantitative PCR was performed as previously described^84^. DNA quantities were normalized by spectrophotometry and %LRT was normalized to WT after subtracting Cq values from matched EFV-treated samples.

For immunoblotting, equal volumes of viral lysates separated by electrophoresis on Bolt 4–12% Bis-Tris Plus gels (Life Technologies) were transferred using Trans-Blo Turbo Mini PVDF Transfer Packs (Bio-Rad). Membranes blocked for 1 h at room temperature were probed overnight at 4 °C in 5% non-fat dry milk with 1:2,000 anti-CA (Abcam ab63917) or 1:5,000 anti-IN^85^. The following day, immunoblots were probed using goat anti-rabbit secondary antibody conjugated to horseradish peroxidase (Agilent P0448), treated with enhanced chemiluminescence detection reagents (Cytiva), and imaged on ChemiDoc Imaging System (Bio-Rad). Adjusted relative band intensity values for anti-IN immunoblots were determined using ImageJ software with anti-CA bands acting as loading controls. A vertical rectangular selection was made over each lane and plotted as a histogram for both immunoblots. For anti-CA, p24 and p25 bands were combined. The area under the curve representing the protein of interest was calculated using the "wand" tracing tool. A "percentage value" was assigned to each sample by dividing the area of each individual peak by the sum of all peaks. Relative band intensity to WT was calculated for each immunoblot. Finally, adjusted relative density was determined by dividing anti-IN relative density by respective anti-CA relative density.

For TEM, virus pellets were resuspended in 1 mL fixative solution (2.5% glutaraldehyde, 1.25% paraformaldehyde, 0.03% picric acid, 0.1 M sodium cacodylate, pH 7.4) and incubated at 4 °C overnight. The preparation and sectioning of fixed virus pellets was performed at Harvard Medical School Electron Microscopy core facility as previously described^86^. Sections (50 nm) were imaged using a JEOL 1200EX transmission electron microscope operated at 80 kV. All images were taken at 30,000X magnification. Micrographs were uploaded to ImageJ and analyzed using the Cell Counter plugin, which allows numbers to be assigned to individual virions. Categories "Mature", "Immature", "Empty", and "Eccentric" were assigned based on previously described IN mutant phenotypes^66^.

## Acknowledgements

Funding: We are grateful for funding from the following sources: NIH grants U01 AI136680 awarded to D.L. and M.K., R01 AI146017 awarded to D.L., U54 AI170855 awarded to D.L, M.K., and R.C., R01 AI184419 to D.L. and M.K., R37 AI039394 and U54 AI170791 awarded to A.N.E., the Intramural Program of the National Institute of Diabetes and Digestive Diseases, NIH to R.C. D.L. is also grateful to the Margaret T. Morris Foundation and the Hearst Foundations, NIH-NCI CCSG P30 CA014195, and R01 GM151305 for funding support. T.J. acknowledges support by the Pioneer Fellowship from the Salk Institute. A.B. acknowledges support from the Schmidt AI Postdoctoral Fellowship from the Eric and Wendy Schmidt Foundation. T.S.S. acknowledges support from an F32 fellowship from the NIH GM148049-03. The molecular graphics and analyses were performed with the USCF Chimera package (supported by NIH P41 GM103311). The cryo-EM data for this project was collected at Scripps Research with equipment supported by NIH grant S10OD032467.

## Author Contributions

T.J. with help from Z.S. produced IN and IN-IBD complexes; Z.S. with help from T.J. performed cryo-EM experiments. T.J. and Z.S. processed cryo-EM datasets. Z.S. built and refined the IN-IBD model; B.Z. designed early IN clones; T.D., H.J.S., G.G., and T.J. purified IN mutant proteins; T.D. performed RNA bridging and SEC experiments; A.B. and Y.Z. assisted with image processing of IN-IBD datasets; S.J., J.G, and Z.L. generated viral clones and performed virology experiments; M.L. purified the WT HIV intasome; D.P. collected the WT HIV intasome cryo-EM dataset and performed initial analyses; Z.Z. analyzed the HIV intasome cryo-EM dataset, refined individual halves, and built the composite map; D.L., T.J., and Z.Z. built the intasome hexadecamer model; T.S.S. developed and supported the computation environment for data processing; S.A. and L.A. developed the graphene protocol for cryo-EM; R.C., A.N.E., M.K., and D.L. oversaw experiments and funding support. D.L. and T.J. wrote the manuscript, with help from A.N.E., M.K., as well as other authors for individual sections.

## Data and materials availability

All data needed to evaluate the conclusions in the paper are present in the paper and/or the Supplementary Materials. The final cryo-EM reconstructions were deposited with the EMDB under accession codes EMD-44962 (IN-IBD tetramer), EMD-70530 (IN tetramer), EMD-45151 (composite map of the hexadecameric HIV intasome), EMD-45103 (consensus map of the intasome), EMD-45104 (top half), and EMD-45150 (bottom half), and the fitted coordinates with the PDB under accession codes 9BW9 (IN-IBD tetramer), and 9C29 (poly-Ala model of the hexadecameric HIV intasome).

---

**Supplementary Figure 1.**
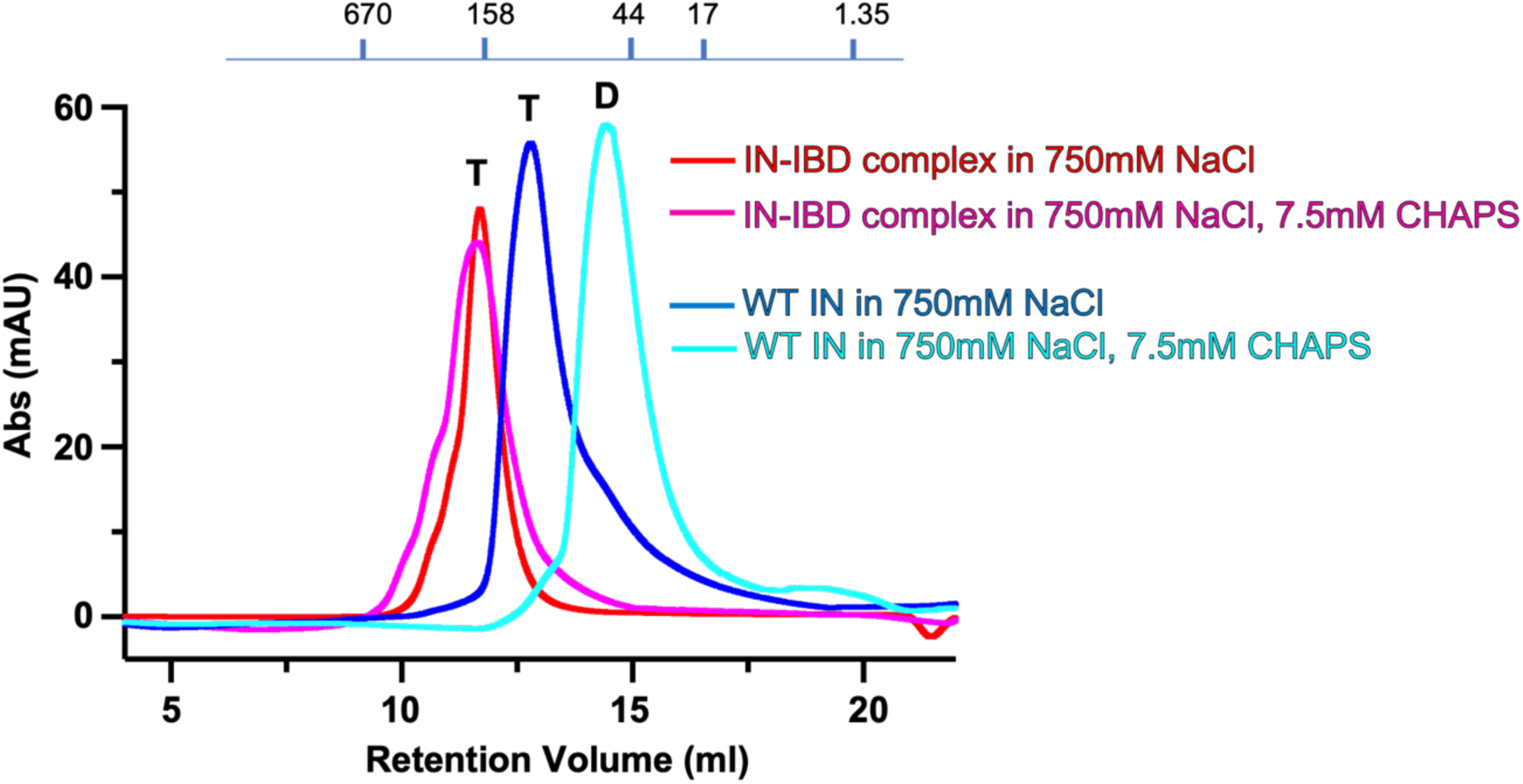
SEC analysis of WT IN and IN-IBD complexes in two buffer conditions. The IN-IBD complex or IN alone was eluted in buffers containing 750 mM NaCl in the presence or absence of CHAPS. In the absence of the IBD, the multimeric state of IN changed as a function of CHAPS, but the IN- IBD complex eluted as a tetramer irrespective of CHAPS in the buffer. T: tetramer, D: dimer. The migration positions of protein standards are indicated atop the chromatograms.

**Supplementary Figure 2.**
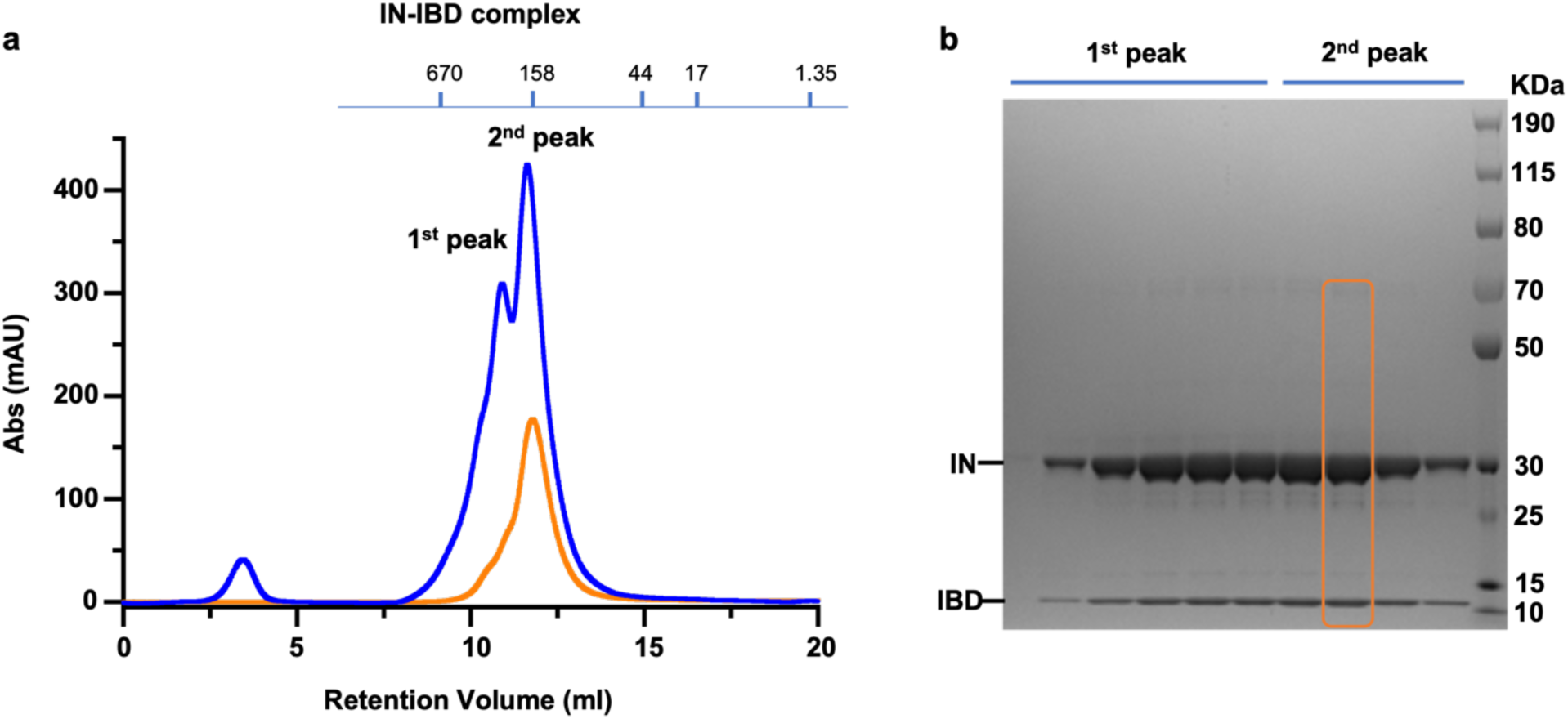
Purification of the IN-IBD complex. (**a**) The IN-IBD complex was initially purified and eluted from SEC, represented by the blue trace. The peak fraction from the 2^nd^ peak of the blue trace was re-injected into the same gel filtration column and eluted again (orange trace), yielding one peak and indicating that a single round of SEC is sufficient to select the tetrameric species. (**b**) SDS-PAGE analysis of the individual fractions of the initial SEC run (blue trace). The boxed fraction was selected for cryo-EM.

**Supplementary Figure 3.**
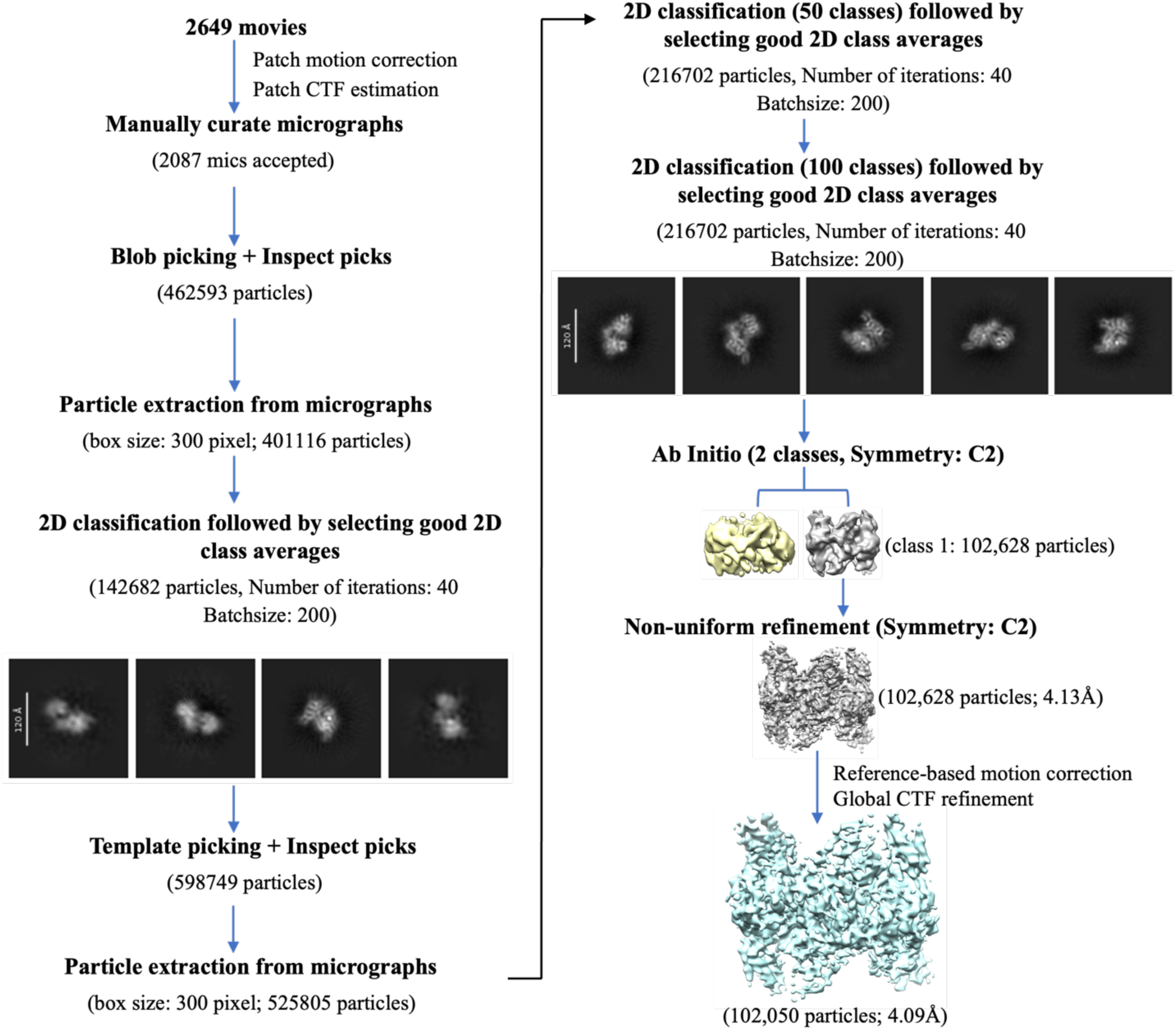
Cryo-EM data-processing workflow for the IN-IBD complex. Schematic of pre-processing, particle selection, classification, and refinement procedures used to generate the 4.1 Å map in this study (see Methods for details).

**Supplementary Figure 4.**
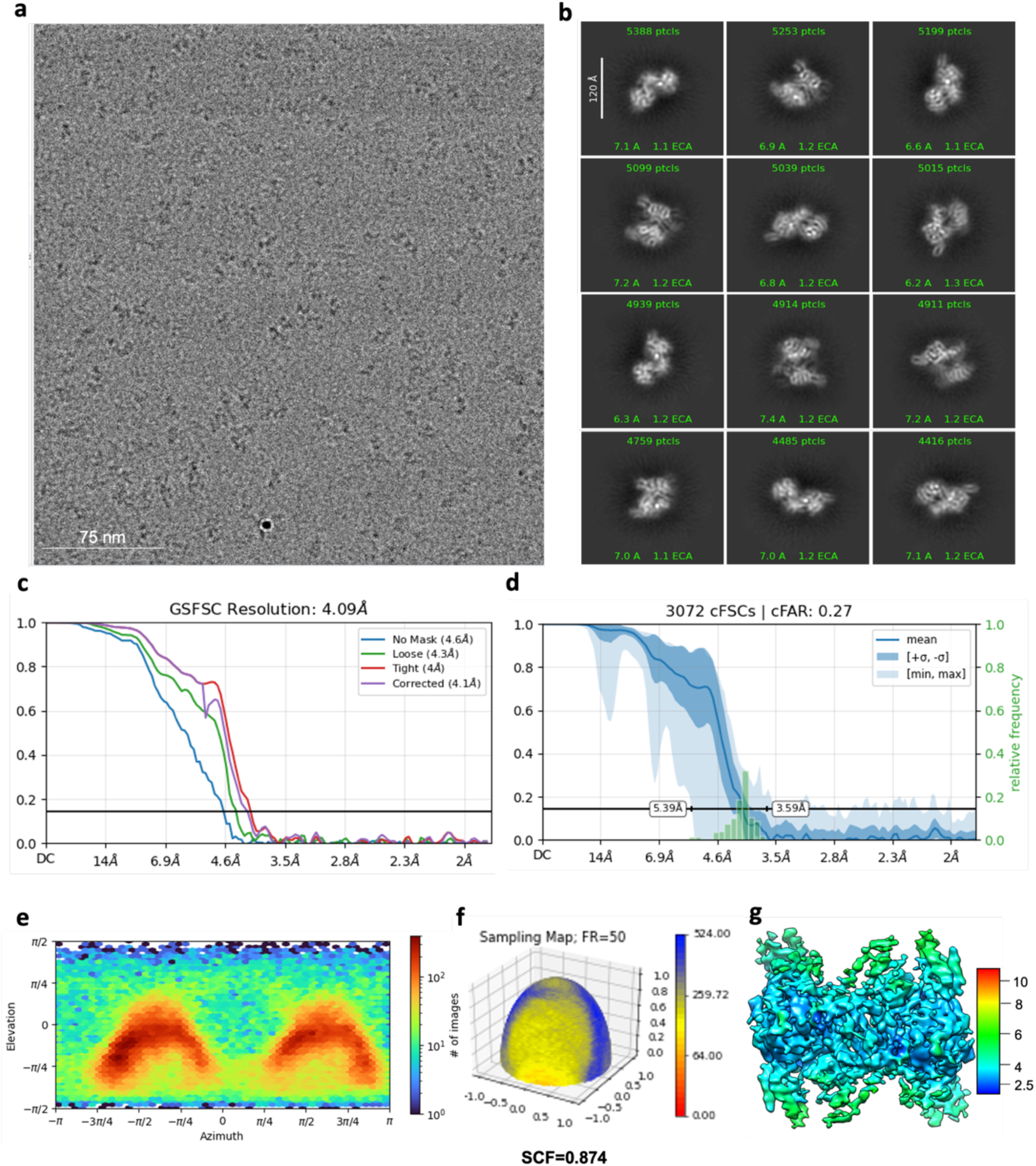
Cryo-EM validations for the IN-IBD tetramer. (**a**) Representative cryo-EM micrograph of IN-IBD complexes. (**b**) Representative 2D class averages of IN-IBD complexes. (**c**) Fourier shell correlation (FSC) curves derived from two half-maps with FSC cutoffs at the 0.143 threshold indicated. (**d**) Directional 3D conical Fourier Shell Correlation (cFSC) plot from non-uniform refinement. (**e**) Euler angle orientation distribution plot from the non-uniform refinement. (**f**) Surface sampling plot for the Euler angle distribution, with the sampling compensation factor (SCF) value indicated below. (**g**) Cryo-EM map colored by local resolution.

**Supplementary Figure 5.**
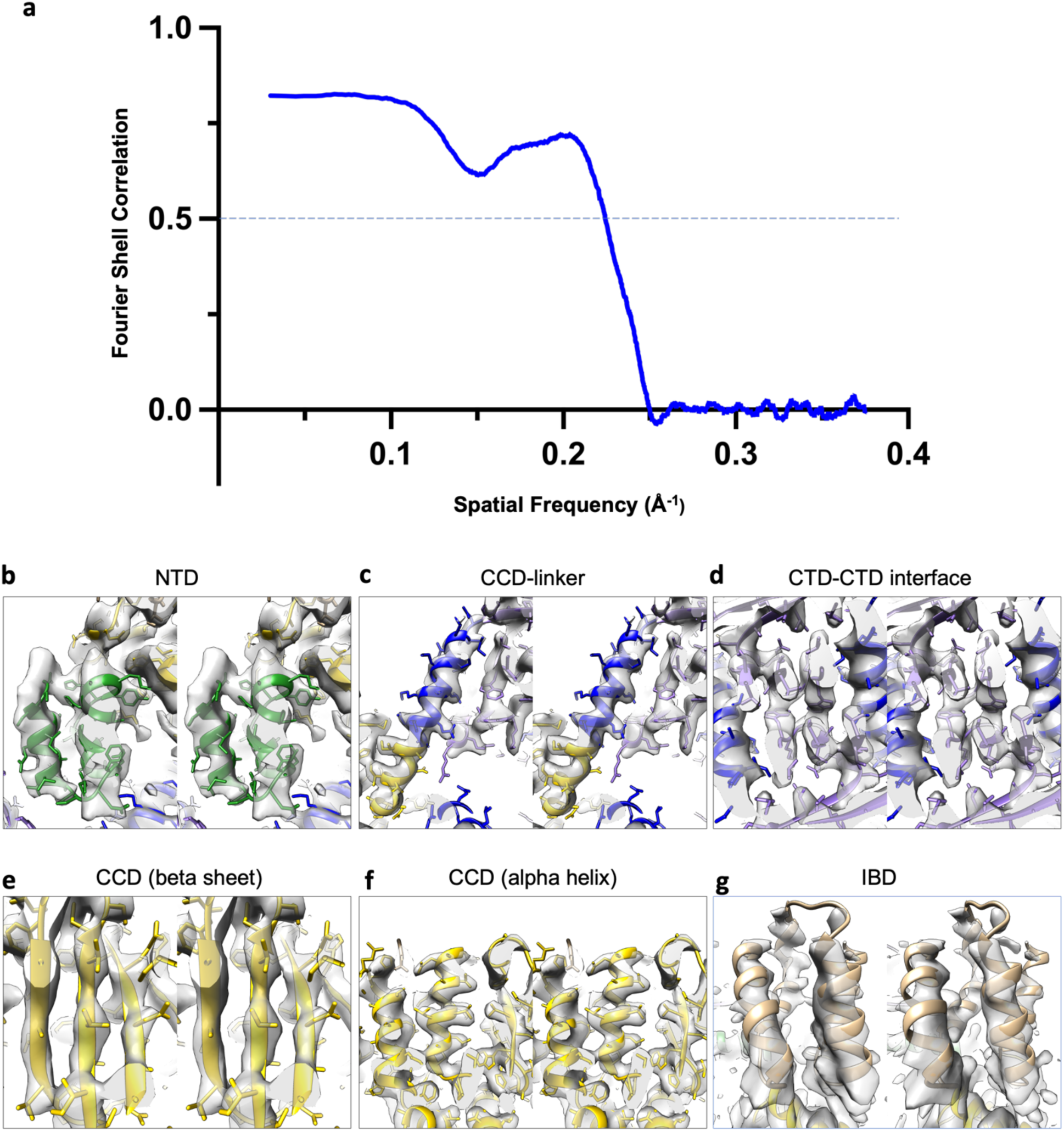
Validation for IN-IBD atomic model. (**a**) FSC curve comparing the full map with the atomic model derived from Phenix real space refinement, with FSC cutoff at the 0.5 threshold indicated. (**b-g**). Stereo views of the atomic model fitted into the cryo-EM density map across different regions of the IN-IBD complex. The experimental cryo-EM density is shown in gray. The atomic model is color-coded by domains in accordance with the other figures. Different views correspond to (**b**) the NTD (green), (**c**) the region between the CCD (yellow) and the CCD-CTD linker (blue), (**d**) the two-fold CTD– CTD interface (purple), including regions of the linkers, (**e**) a β-sheet region within the CCD, (**f**) several α- helical regions within the CCD, and (**g**) the IBD (brown).

**Supplementary Figure 6.**
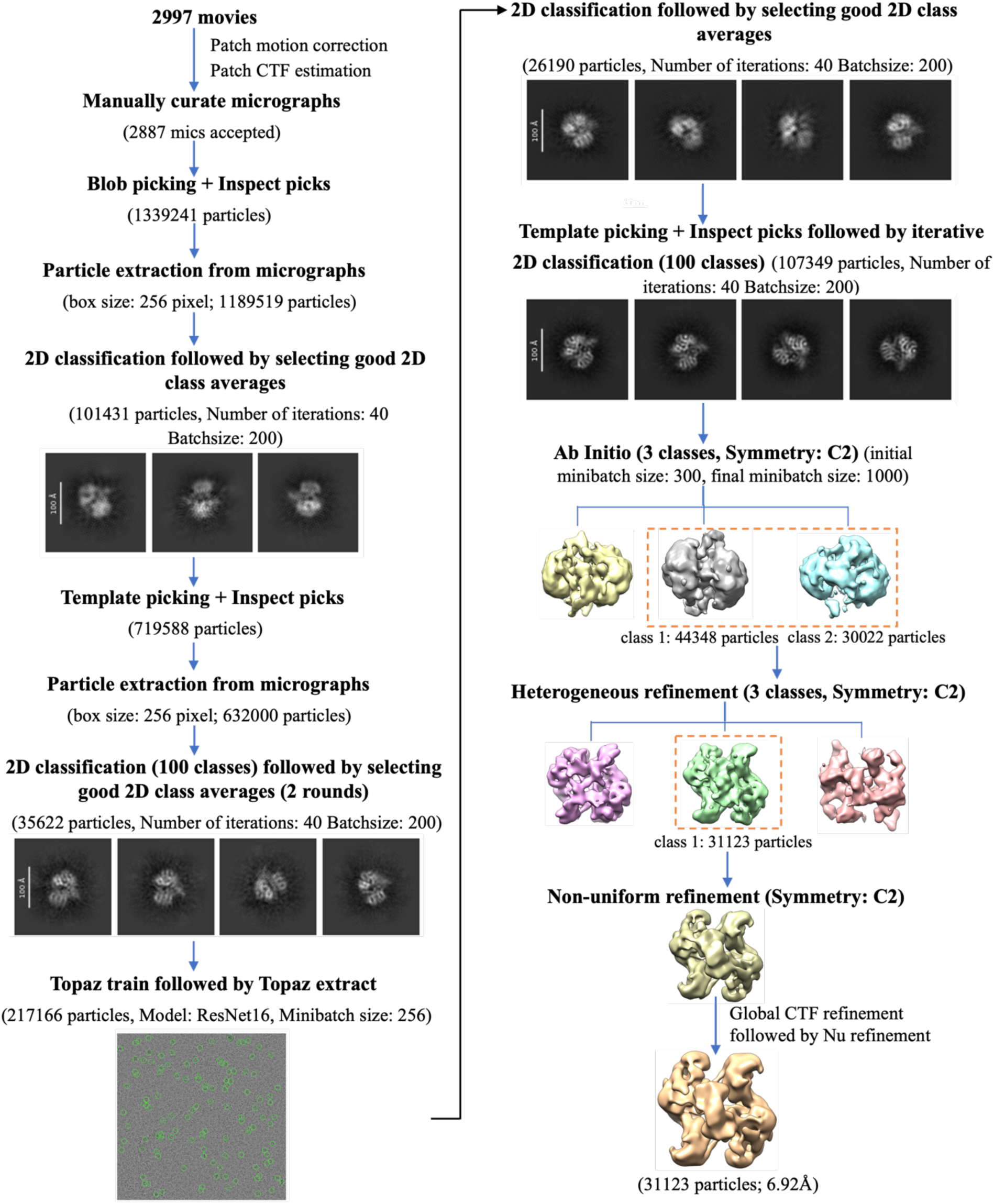
Cryo-EM data-processing workflow for the apo IN tetramer complex. Schematic of pre-processing, particle selection, classification, and refinement procedures used to generate the 7 Å map in this study (see Methods for details).

**Supplementary Figure 7.**
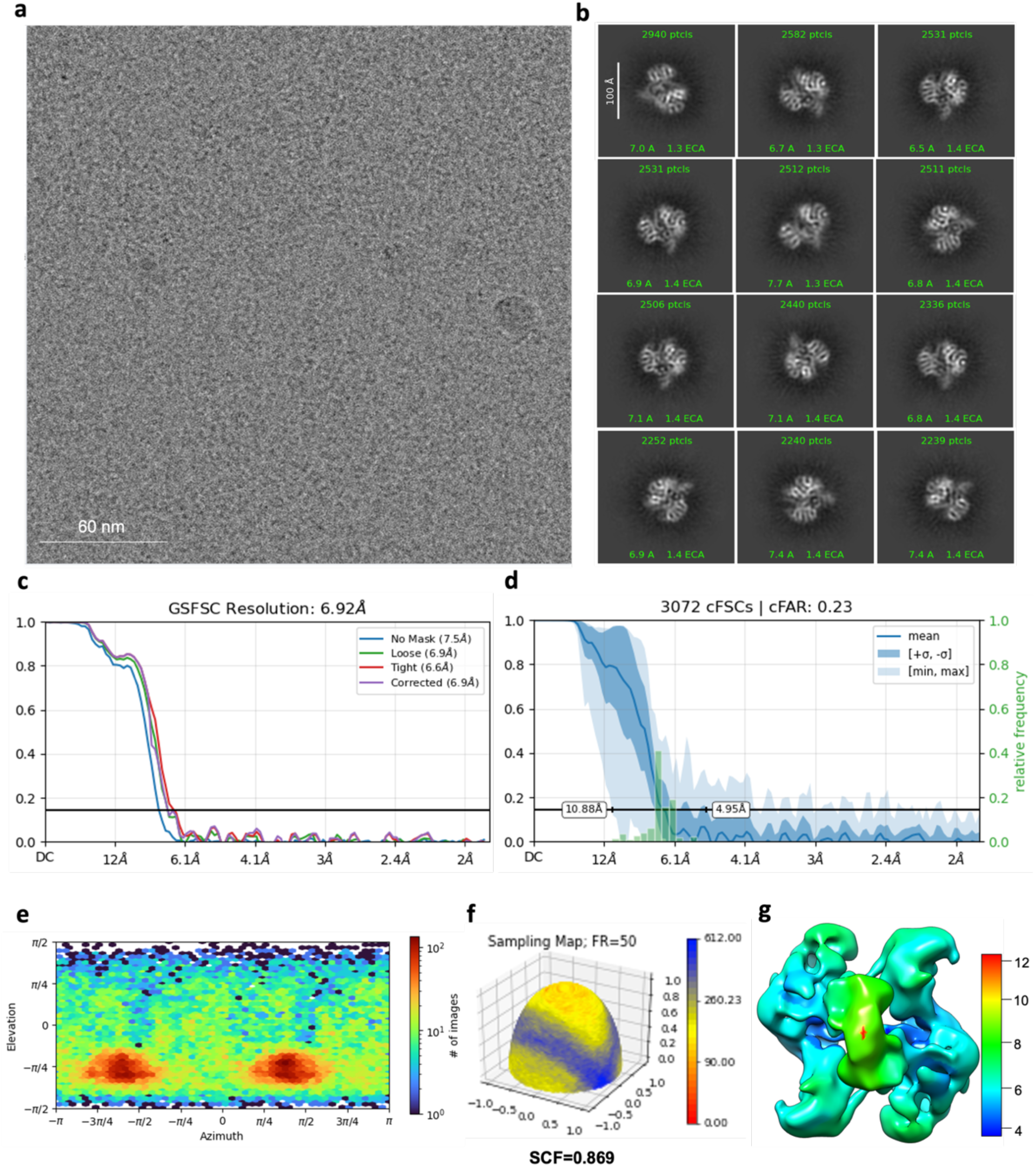
Cryo-EM validations for the apo IN tetramer. (**a**) Representative cryo-EM micrograph of apo IN complexes. (**b**) Representative 2D class averages of the apo IN complexes. (**c**) Fourier shell correlation (FSC) curves derived from two half-maps with FSC cutoffs at the 0.143 threshold indicated. (**d**) Directional 3D conical Fourier Shell Correlation (cFSC) plot from non-uniform refinement. (**e**) Euler angle orientation distribution plot from the non-uniform refinement. (**f**) Surface sampling plot for the Euler angle distribution, with the sampling compensation factor (SCF) value indicated below. (**g**) Cryo-EM map colored by local resolution.

**Supplementary Figure 8.**
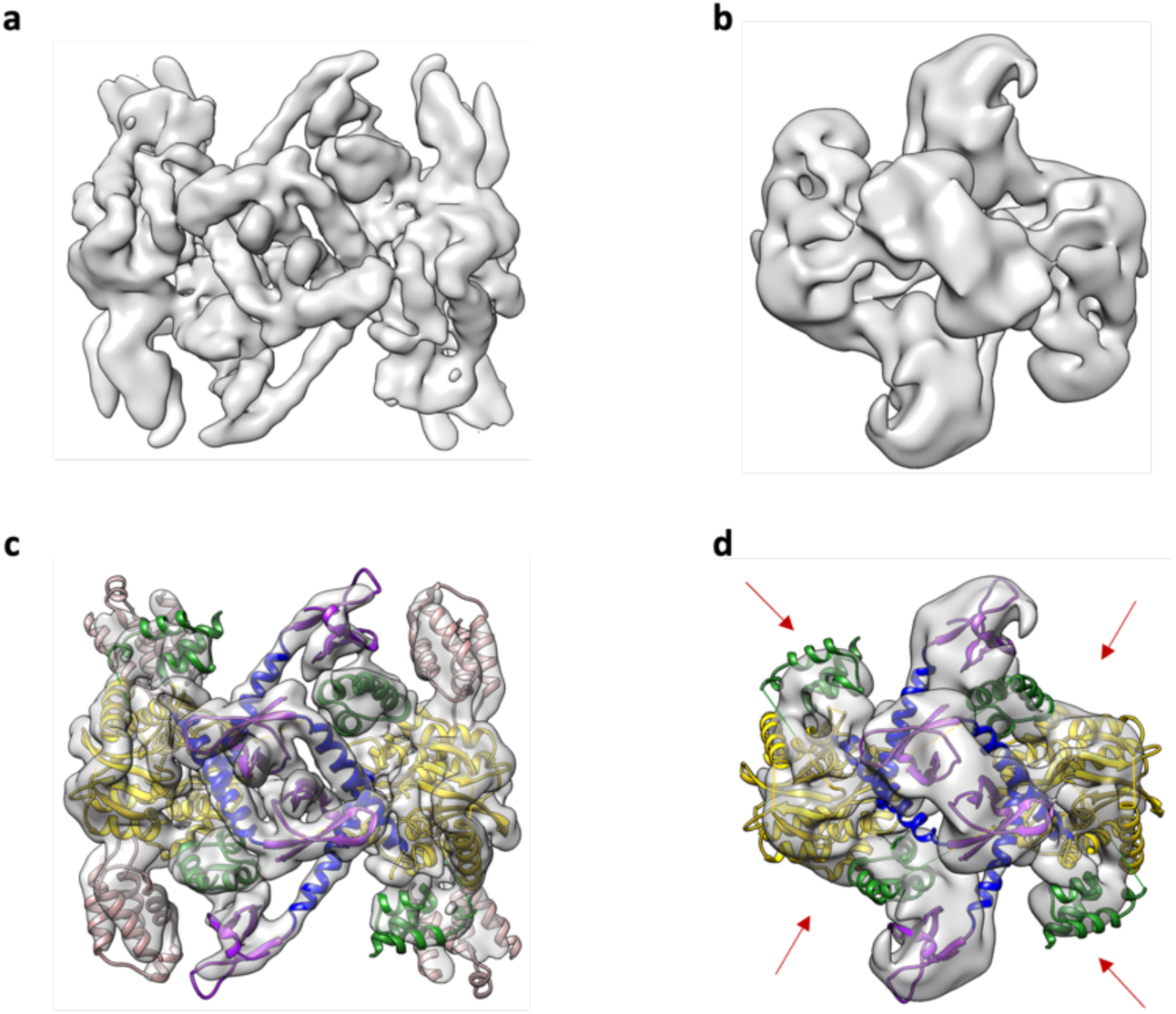
**Comparison of IN-IBD and apo IN cryo-EM reconstructions**. (**a**) Unsharpened IN-IBD map and (**b**) unsharpened apo IN map, displayed in the same relative orientation along the two-fold symmetry axis. (**c**) Atomic model of IN-IBD docked into IN-IBD density map. (**d**) the same atomic model of the IN-IBD docked into the apo IN map. For clarity, the four IBDs were removed from the model and the red arrows indicate the missing IBD positions. The IN-IBD model is colored by domain as in Figure 1.

**Supplementary Figure 9.**
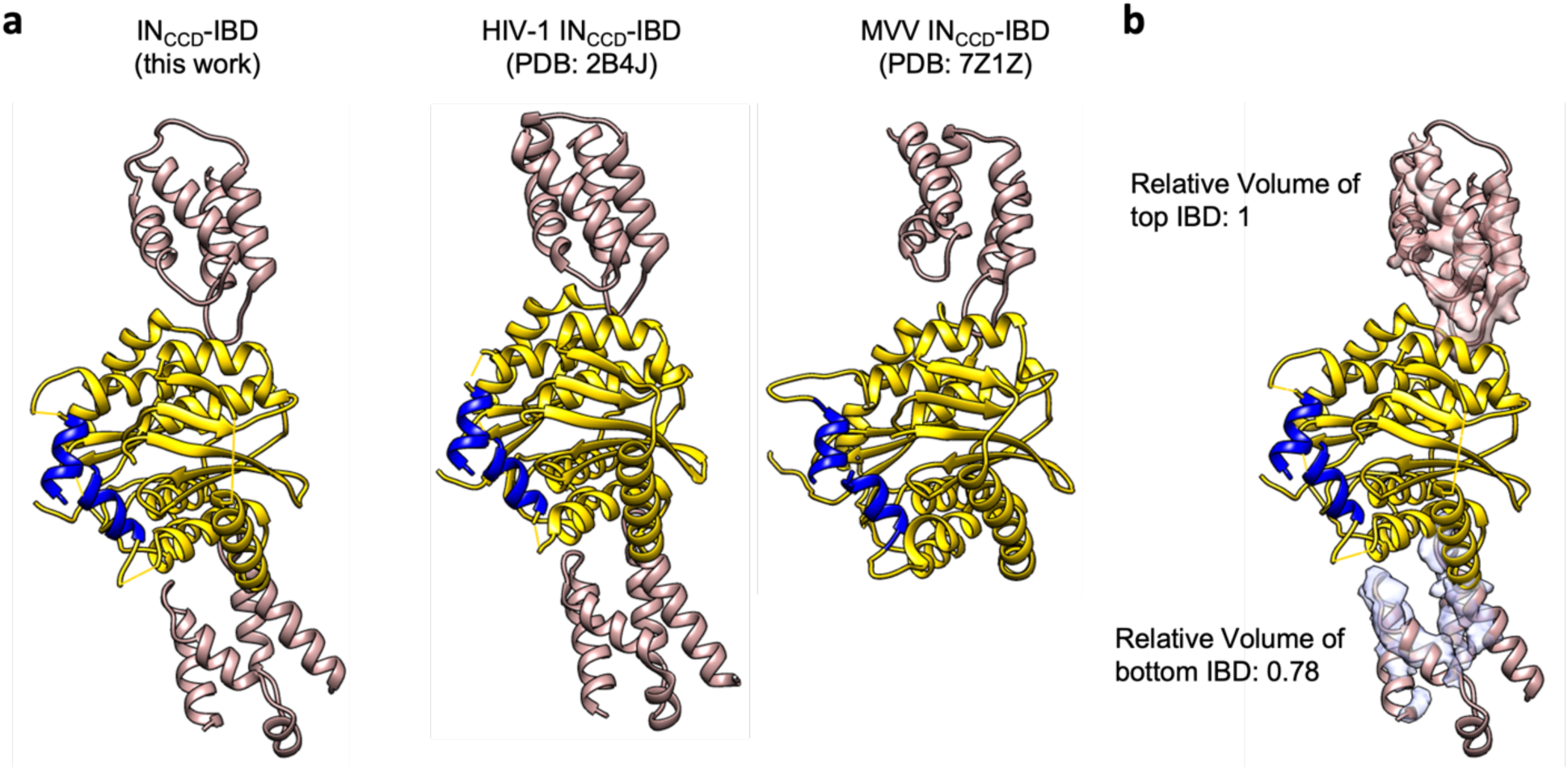
Comparison of IBD organization within the IN tetramer with prior structures. (**a**) Left panel: an individual IN-IBD dimer extracted from the full-length IN-IBD tetramer. Middle panel: IN-IBD dimer from crystalized HIV-1 IN_CCD_ dimer complexed to IBD (PDB 2B4J)^53^. Right panel: one IBD bound to the IN_CCD_ dimer from the conserved intasome core of the MVV intasome (PDB 7Z1Z)^9^. (**b**) An individual IN-IBD dimer is extracted from the full-length IN-IBD tetramer (as in panel **a**), with the two segmented cryo-EM densities for the IBDs displayed in transparency. The enclosed volume of top and bottom IBD densities are calculated respectively in Chimera, and the relative volumes are labeled next to the density map. The CCD dimers are colored in yellow and IBD are in brown. The beginning of the CCD-CTD linker is colored blue at residue 196 for HIV-1 or its analogue in MVV. For clarity, NTDs and CTDs were omitted.

**Supplementary Figure 10.**
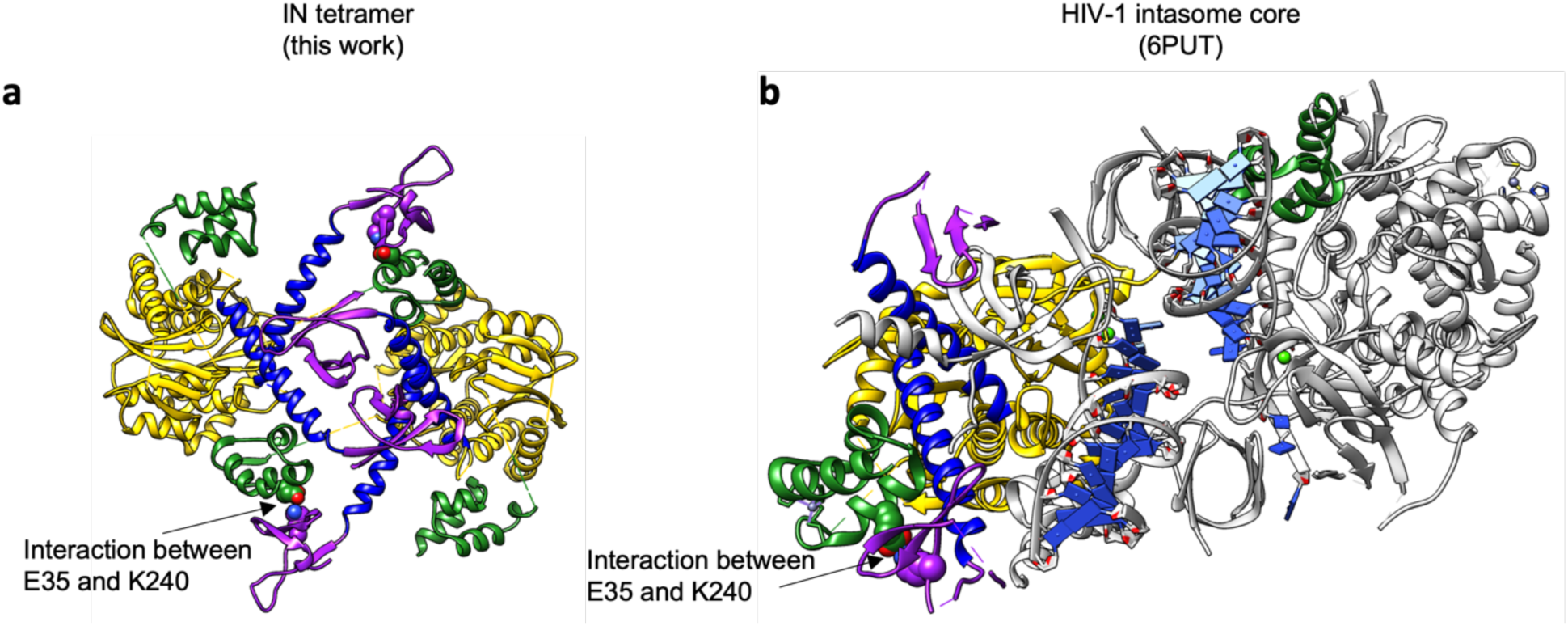
**Comparison of the CTD:NTD interface between the IN tetramer and the HIV-1 intasome**. (**a**) CTD-NTD interface in the IN tetramer showing interaction between E35 and K240. (**b**) CTD-NTD interface in the conserved core of the HIV-1 intasome (PDB 6PUT)^12^ showing interaction between E35 and K240. The colors are as in Figure 1c-d (NTD, green; CCD, yellow; CCD-CTD, blue; CTD, purple). Other regions of the HIV-1 conserved intasome core are shown in gray.

**Supplementary Figure 11.**
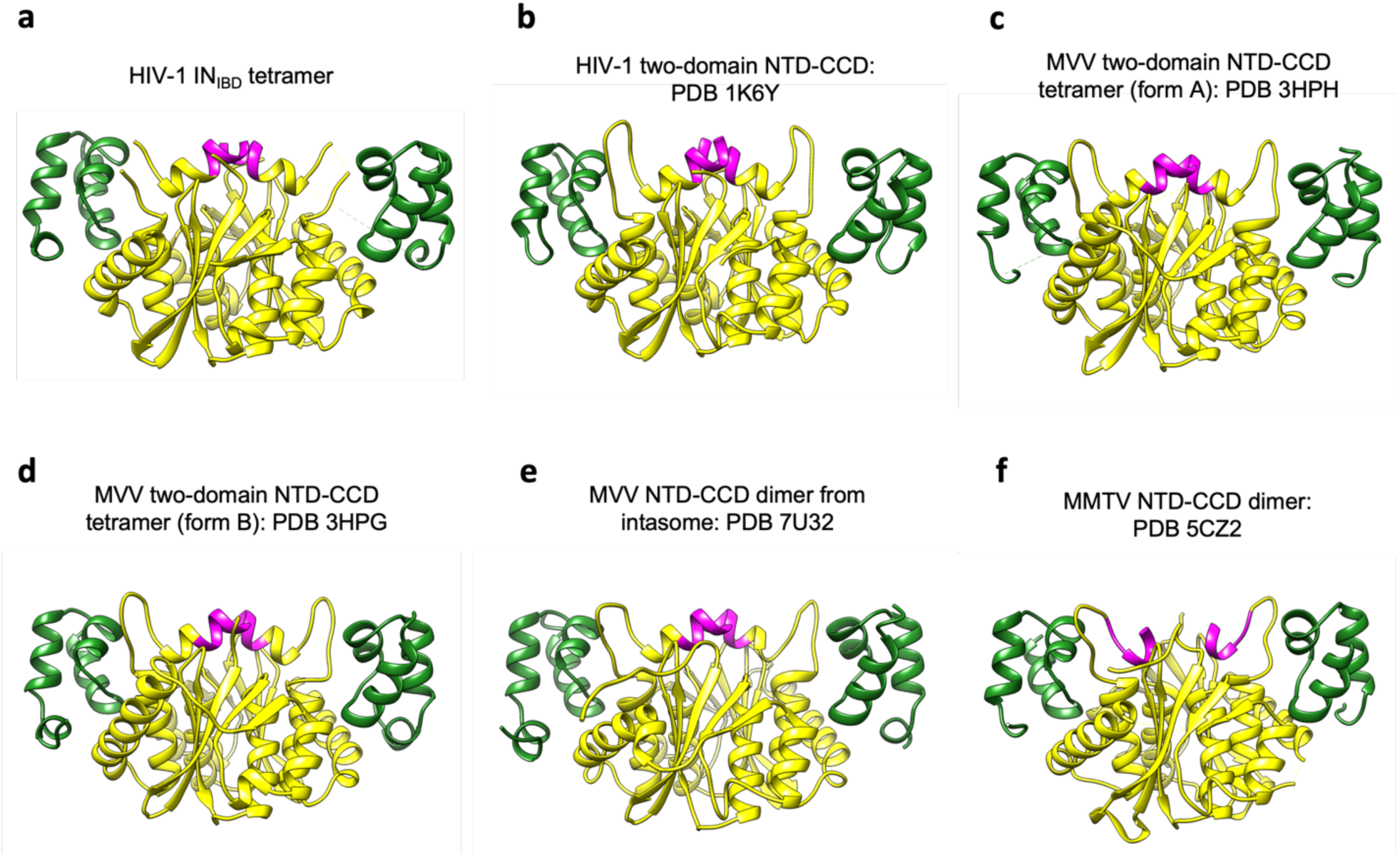
**Comparison of NTD and CCD organization between the IN tetramer and prior structures**. (**a**) NTD-CCD dimer from HIV-1 full-length IN tetramer. (**b**) NTD-CCD dimer from HIV-1 NTD-CCD two-domain construct, which crystalized in a tetrameric form (PDB 1K6Y)^52^. (**c-d**) NTD-CCD dimer from two crystallographic forms of MVV IN tetramers, PDB 3HPH and 3HPG, respectively. (**e**) NTD-CCD dimer selected from an MVV intasome solved by cryo-EM (PDB 7U32). The three other unique dimers within the intasome assembly similarly had density occupying the NTD:CCD interface. (**f**) Crystal structure of mouse mammary tumor virus (MMTV) IN NTD-CCD construct (PDB 5CZ2). In all cases, the NTD and CCD domains are colored by forest green and yellow, respectively. Magenta, CCD-CTD linker regions. All structures are presented in the same orientation. For clarity, the CTD domains in panels **a** and **e** were omitted.

**Supplementary Figure 12.**
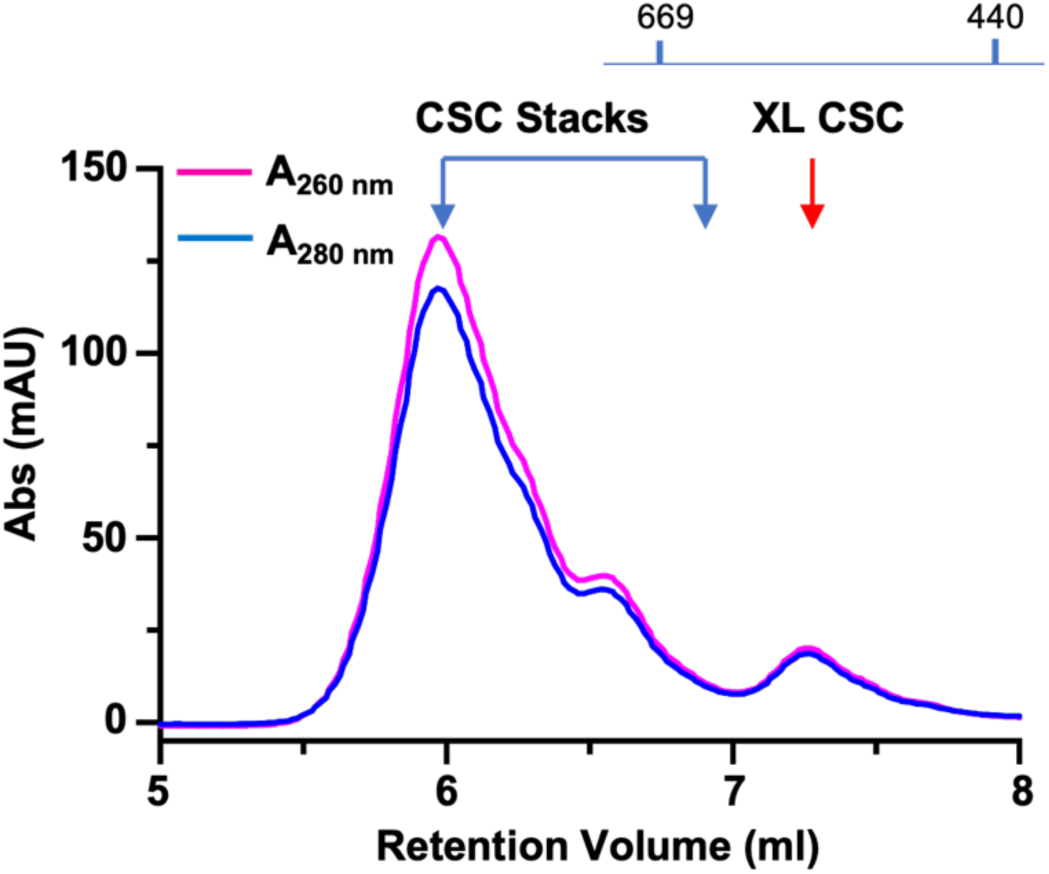
Purification of the crosslinked WT HIV-1 intasome. Crosslinked intasomes were purified by gel filtration on a TSKgel UltraSW HPLC column in 20 mM Tris pH 6.2, 0.5 mM TCEP, 1.0 M NaCl, 5.0 mM MgCl_2_, and 10% (w/v) glycerol. The gel filtration profile for the purification is shown. All species eluting earlier than ∼7 mL retention volume correspond to intasome stacks, which have been previously described for HIV-1^12^, SIV^19^, and MVV intasomes^10^. These stacks contain multiple vDNAs copies and are not physiologically relevant. Intasomes used for cryo-EM studies are indicated by the red arrow. Elution volumes for 669 kDa and 440 kDa molecular mass standards are indicated.

**Supplementary Figure 13.**
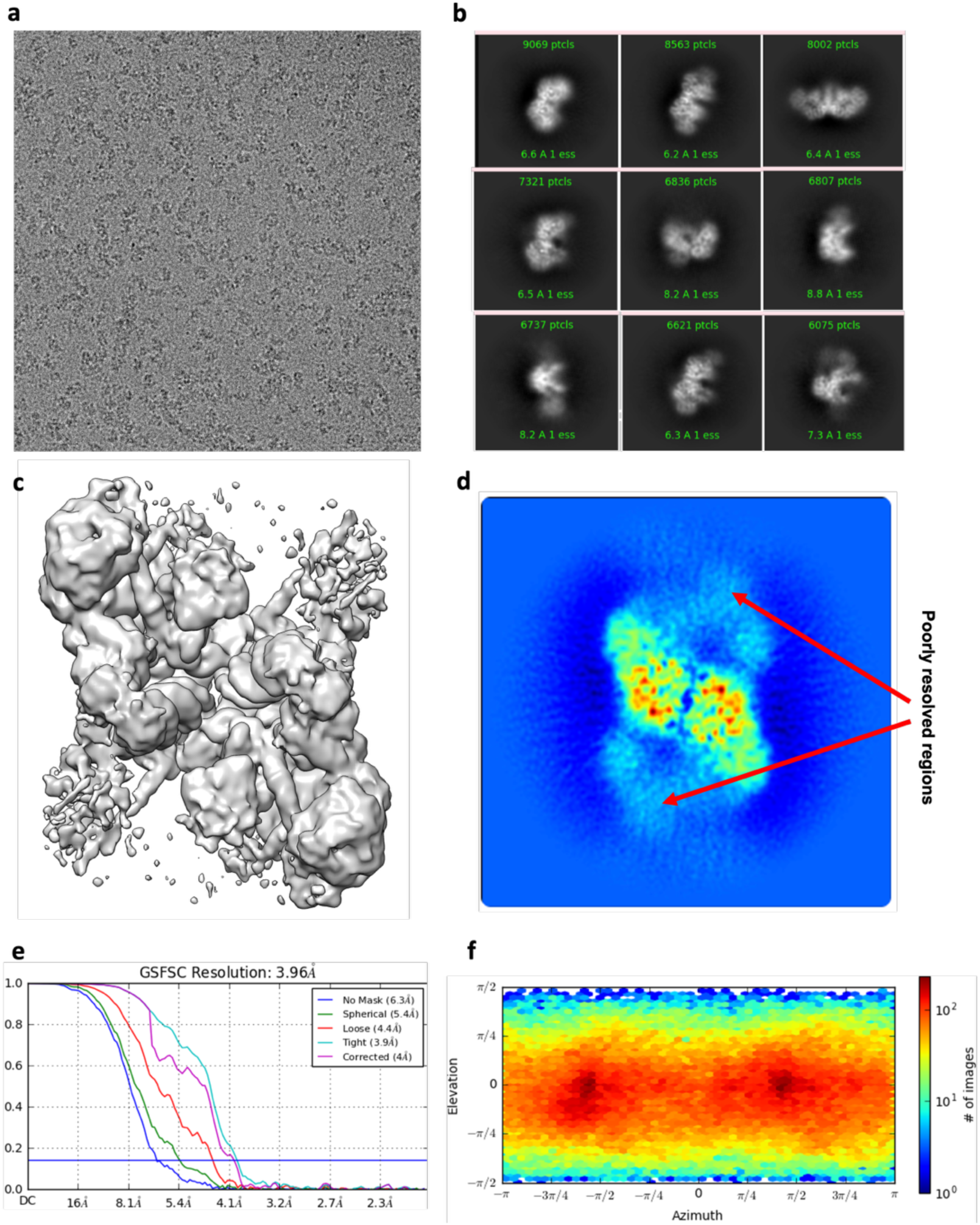
Initial results from cryo-EM processing of the crosslinked HIV-1 intasome. (**a**) Representative micrograph of crosslinked HIV-1 intasomes. (**b**) Representative 2D class averages from the cryo-EM data. (**c**) Cryo-EM map generated in cryoSPARC^49^ using a conventional data processing strategy and after global 3D classification. (**d**) Cross-section of the reconstructed map from 3D classification, showing heterogeneous density for the poorly resolved flanking regions, which correspond to the outer IN subunits. We were unable to recover the missing data using any combination of global classification approaches, with and without symmetry applied. (**e**) FSC curves derived from two half-maps with the 0.143 cutoff threshold indicated. **(f)** Euler angle orientation plot from the cryoSPARC refinement.

**Supplementary Figure 14.**
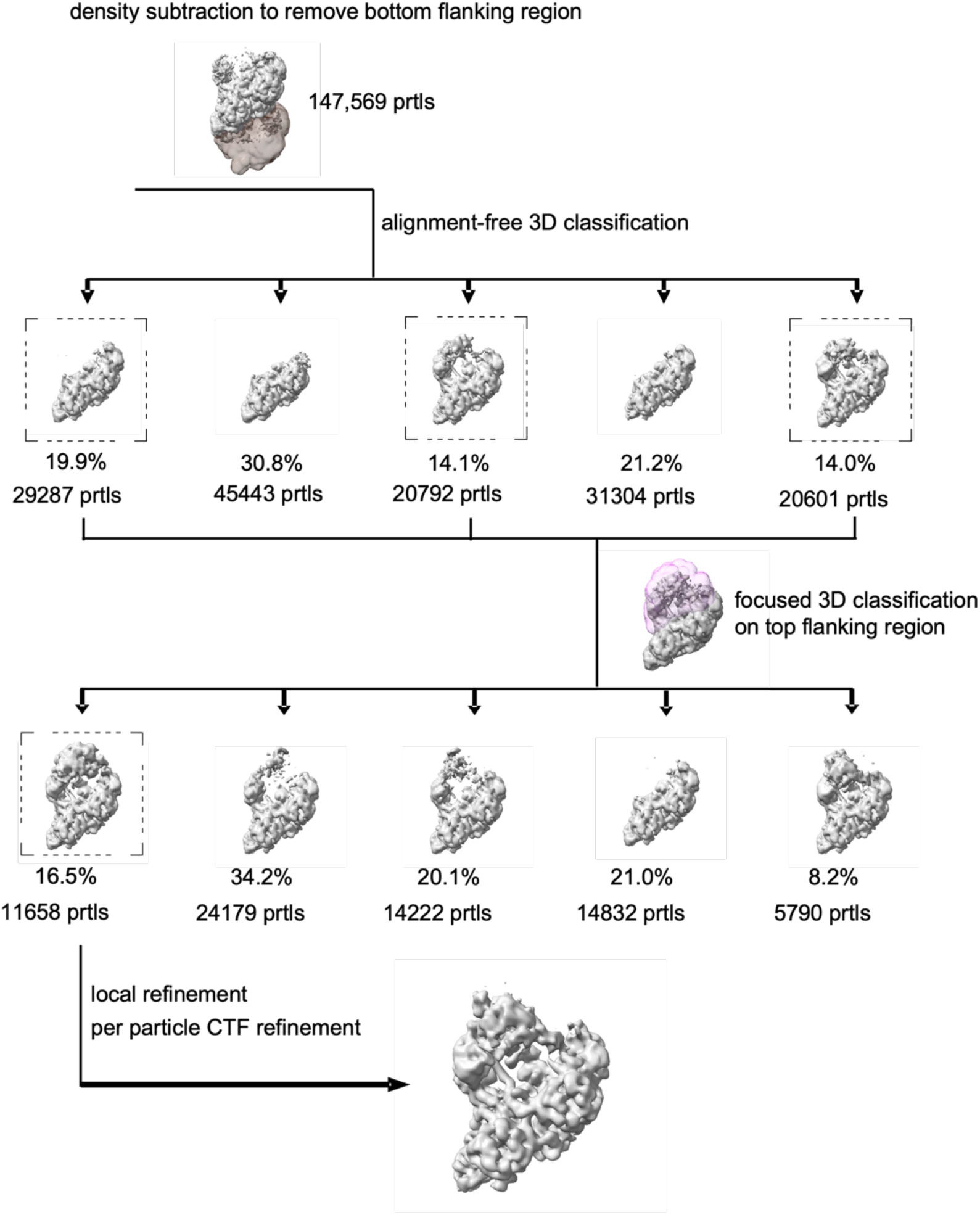
Workflow to resolve the flanking regions within the hexadecameric HIV- 1 intasome. 147,569 particles were subjected to density subtraction to remove signal in the bottom flanking region. The subtracted particles were classified using global 3D classification into five classes. Among them, three classes with potential top flanking densities were selected (dashed box), and the particles within these classes were subsequently subjected to a second round of 3D classification, using a mask focused on the top flanking region. The 11,658 particles from the best class (dashed box) were selected. Finally, we conducted one round of local CTF refinement, followed by a final round of non-uniform refinement^71^, using a mask encompassing the entire volume.

**Supplementary Figure 15.**
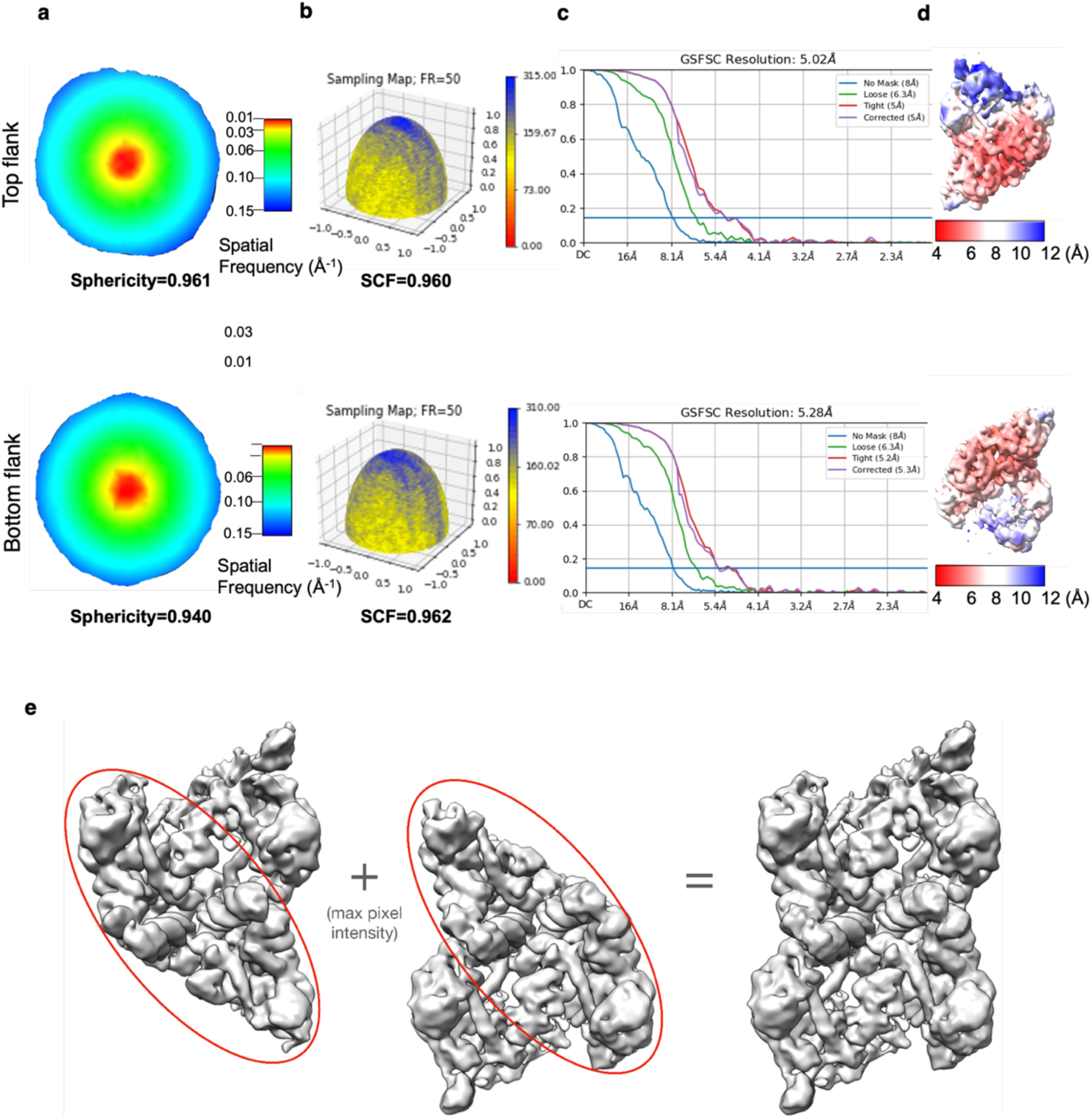
Validation metrics for each independently refined half of the HIV-1 intasome. Evaluation of the two maps containing the top and bottom flanking regions of the hexadecameric intasome. (**a-c**) The top panel corresponds to the map derived from focused classification on the upper flank, and the bottom panel corresponds to the map derived using the same approach on the bottom flank. (**a**) Central slice through the 3DFSC, colored by resolution. (**b**) Surface sampling plot for the Euler angle distribution, with the sampling compensation factor (SCF) value indicated below. (**c**) FSC curves derived from two half-maps with cutoffs at the 0.143 threshold indicated. (**d**) Refined cryo-EM map colored by local resolution. (**e**) The composite map of the HIV-1 intasome hexadecamer was derived from two independently refined halves, summed using maximum pixel intensities for each of the aligned halves.

**Supplementary Figure 16.**
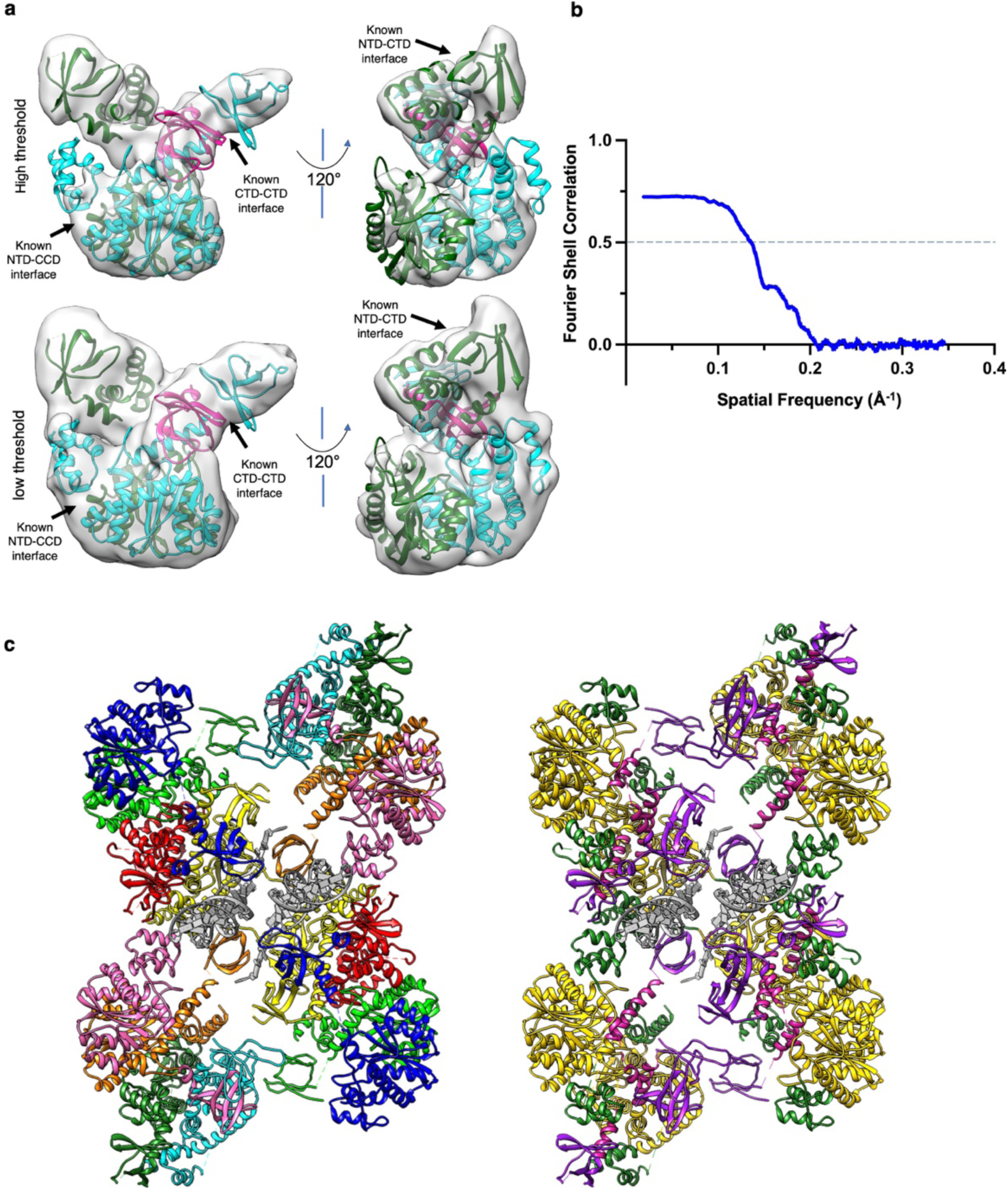
Rigid body docked model of the hexadecameric intasome. (**a**) Comparison of the assembled rigid body model with the experimental cryo-EM density for the poorest regions of the hexadecameric intasome, which includes the complete chains G and H, and the CTD of chain F (as well as the symmetry-mates). These regions correspond to the outer dimeric subunits of the “flanking” IN tetramers, which were not previously observed in HIV-1 intasome cryo-EM maps. Two different docking orientations are shown at high (top) and low (bottom) thresholds. For clarity, the docked models are colored by protomer, as in panel **c** (left image). (**b**) FSC curves comparing the full composite map with the rigid body docked polyalanine model, with FSC cutoff at the 0.5 threshold indicated (**c**) Composite model of the HIV-1 intasome hexadecamer, derived through rigid-body docking of previously determined high-resolution structures of either individual IN domains or multi-domain constructs (see Methods). The model is colored by IN protomer (left) and domain (right).

**Supplementary Figure 17.**
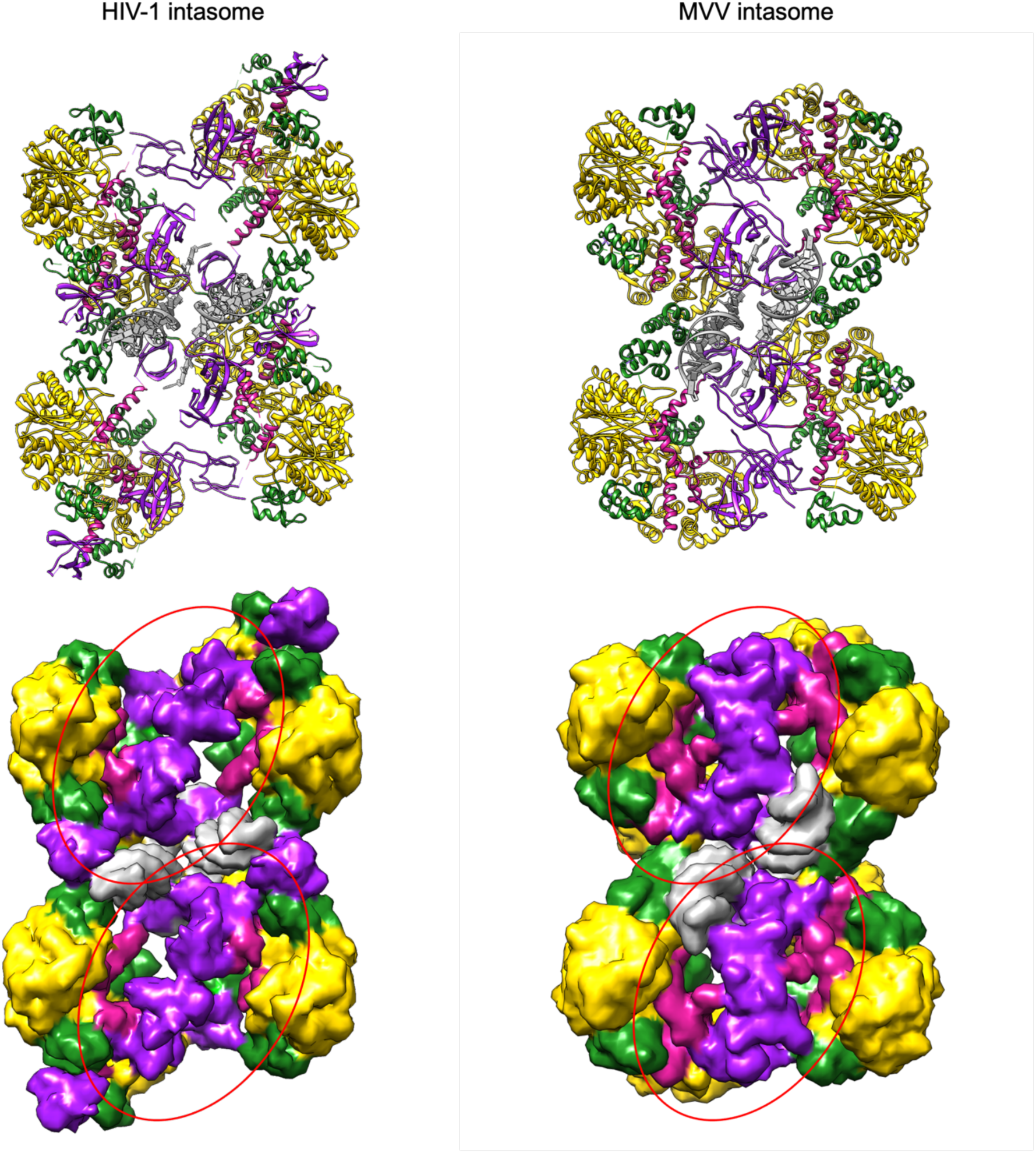
Comparison of the intasome hexadecamers between HIV-1 and MVV. The pseudo-atomic model of the HIV-1 intasome is displayed alongside the atomic model of the MVV intasome (PDB 7U32)^9^. For each model (top), a molecular map (bottom), generated in Chimera at 8 Å resolution, is displayed to show surface views of the assemblies. The models and maps are colored by IN domain. The inter-tetramer CTD-CTD bridge, which has a smaller interface in the HIV-1 intasome compared to the MVV intasome, is circled.

**Supplementary Figure 18.**
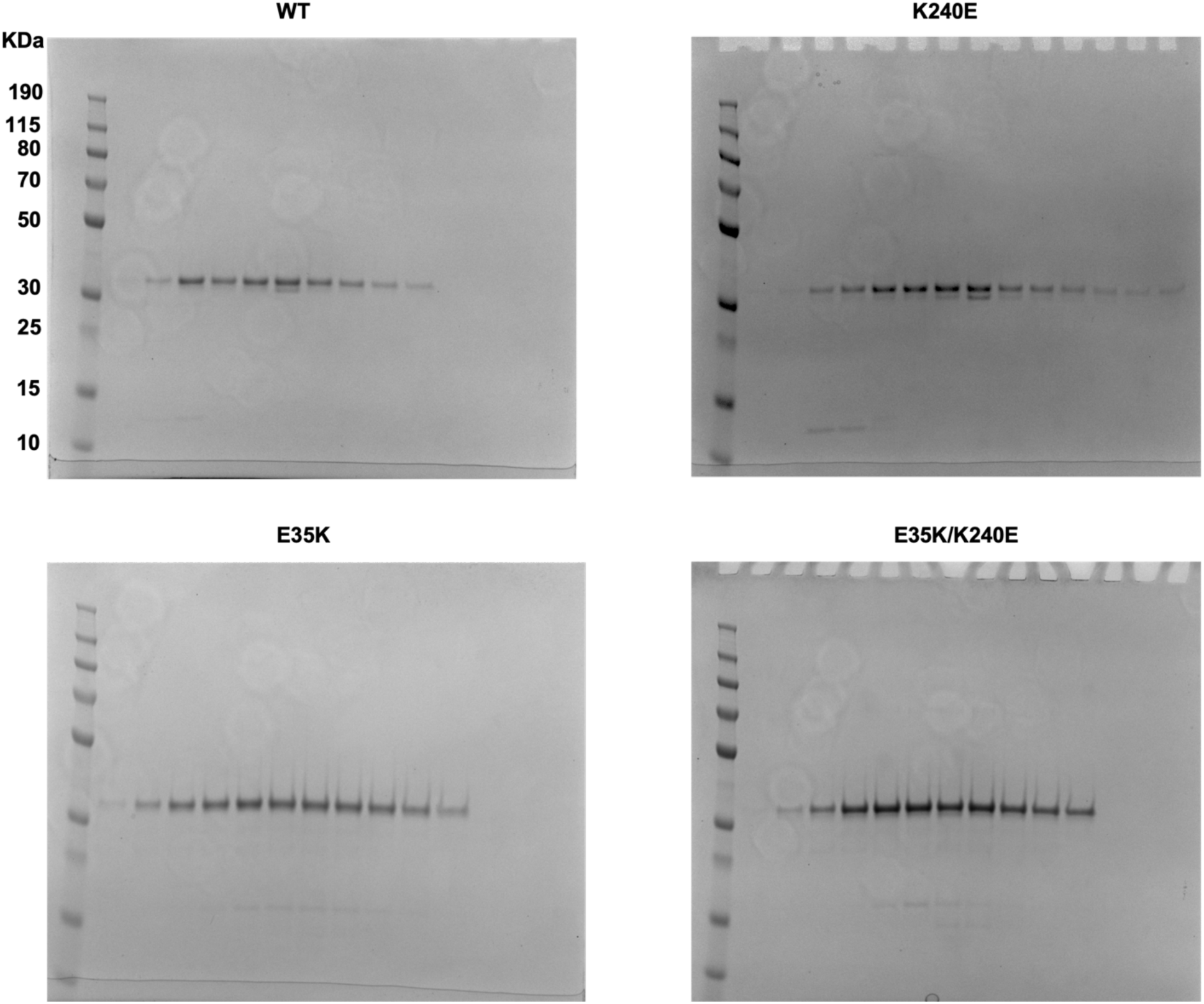
Purification of WT and mutant IN. Purified WT and mutant IN proteins were detected via Coomassie blue staining of SDS-PAGE gels. These gels correspond to mutants tested in Figure 4b-d. Mutants tested in Figure 6e-f shows similar purity.

**Supplementary Table 1.**
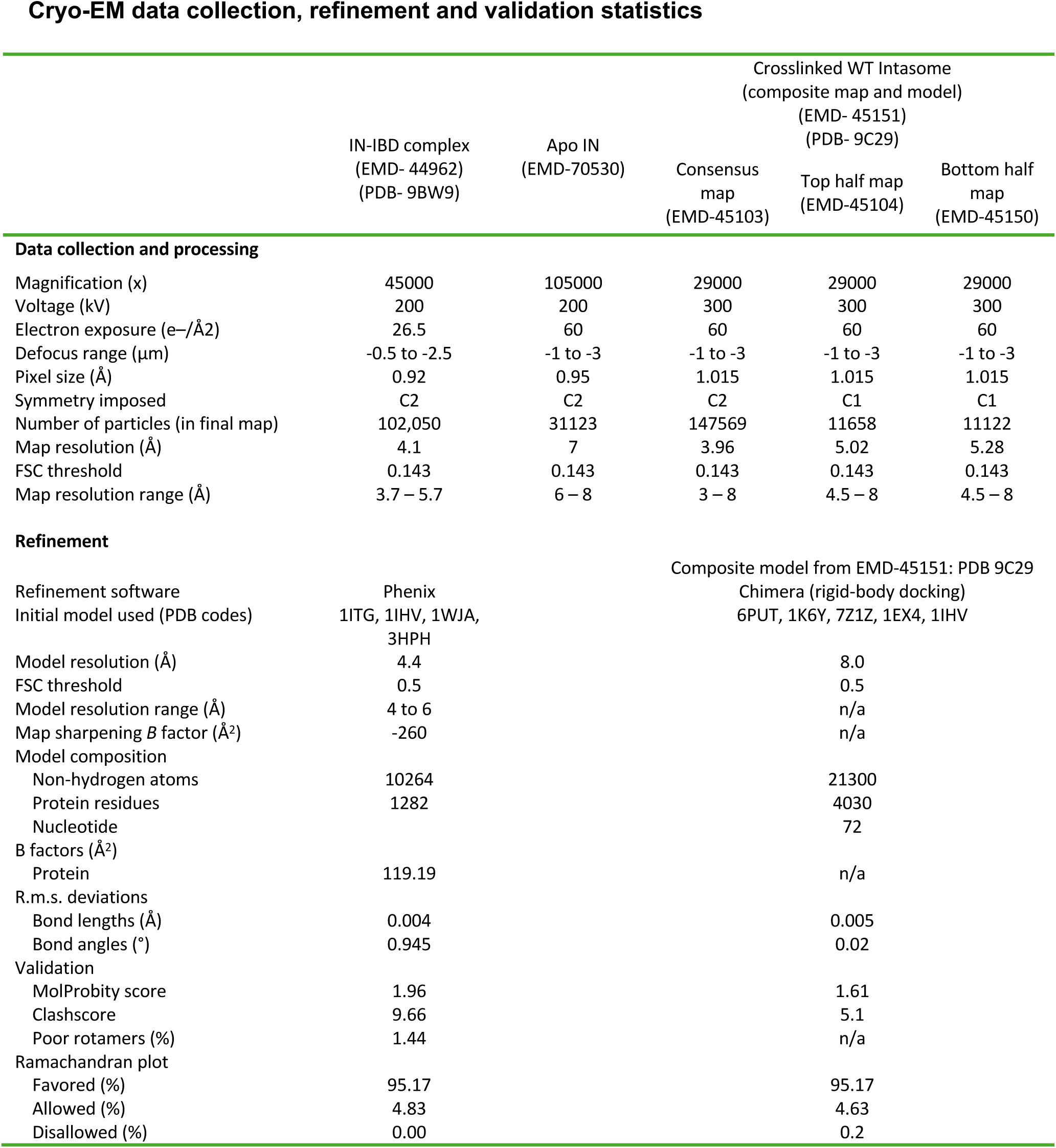
Cryo-EM data collection, refinement, and validation statistics. For the intasome, the crosslinked WT intasome statistics refer to the composite map. The associated EMDB IDs for the consensus map, the top half, and the bottom half are EMD-45103, EMD-45104, and EMD-45150, respectively.

